# Primordial germ cells experience increasing physical confinement and DNA damage during migration in the mouse embryo

**DOI:** 10.1101/2025.03.03.641275

**Authors:** Katharine Goodwin, Theresa Anne Emrich, Sebastian Arnold, Katie McDole

## Abstract

To produce healthy offspring, an organism must pass on its genetic material with high fidelity. In many species, this is accomplished by primordial germ cells (PGCs), which give rise to sperm or eggs. PGCs are often specified far from the future gonads and must migrate through developing tissues to reach them. Failure to do so can result in infertility or germ cell tumours. While PGC migration is well characterized in some species, very little is known about their migration in mammalian embryos. Here, we performed dynamic and quantitative analyses of PGC migration from E7.5 to E9.5 in the mouse embryo, providing the first comprehensive study of the migratory characteristics of PGCs from their point of origin to the gonads. We demonstrate that migrating PGCs are influenced by the surrounding environment and, in contrast to other organisms, extend highly dynamic, actin-rich protrusions to navigate through ECM barriers and tight intercellular spaces. As PGCs migrate through increasingly confined spaces, they undergo significant nuclear deformation and become prone to nuclear rupture and DNA damage. Their migration under confinement may be aided in part by a depleted nuclear lamina that leads to wrinkled nuclear morphology. Our high-resolution and dynamic imaging approaches have uncovered a surprising risk to genome integrity in migrating PGCs, with implications for DNA repair and adaptations in nuclear mechanics in PGCs.

## Introduction

Primordial germ cells (PGCs) are first specified in the proximal posterior epiblast of the mouse embryo at around embryonic day (E) 6.5[1]. Over the next three to five days of development, PGCs migrate through a variety of developing tissues by following external chemotactic signals to reach the sites of the future gonads[2]. PGCs that fail to reach the gonads can undergo apoptosis, leading to infertility, or giving rise to germ cell tumours[3]. PGC migration is evolutionarily conserved among most species, but only to a certain extent[2, 4, 5]. Differences in the size and developmental rate of embryos of different species affects the migratory path of PGCs in terms of distance, duration, and physical obstacles. In the mouse embryo, PGCs first migrate out of the epiblast, through the mesoderm, and into the endoderm on the surface of the embryo by E7.5[6] (Fig. 1a, Supplementary Figure 1a). The involution of the hindgut pocket at the posterior then transports PGCs to the interior of the embryo[7, 8]. From E7.5 to E9.5, PGCs reside in the endoderm while it transforms into the pseudostratified, tubular hindgut endoderm[9, 10]. (Fig. 1a). At E9.5, PGCs then exit the hindgut endoderm and migrate bilaterally through the surrounding mesentery to reach the gonadal ridges[10] (Fig. 1a, Supplementary Figure 1a). While the migratory path of mouse PGCs has been known for some time, the strategies they use to traverse these different tissue environments are less well understood[11].

**Figure 1:**
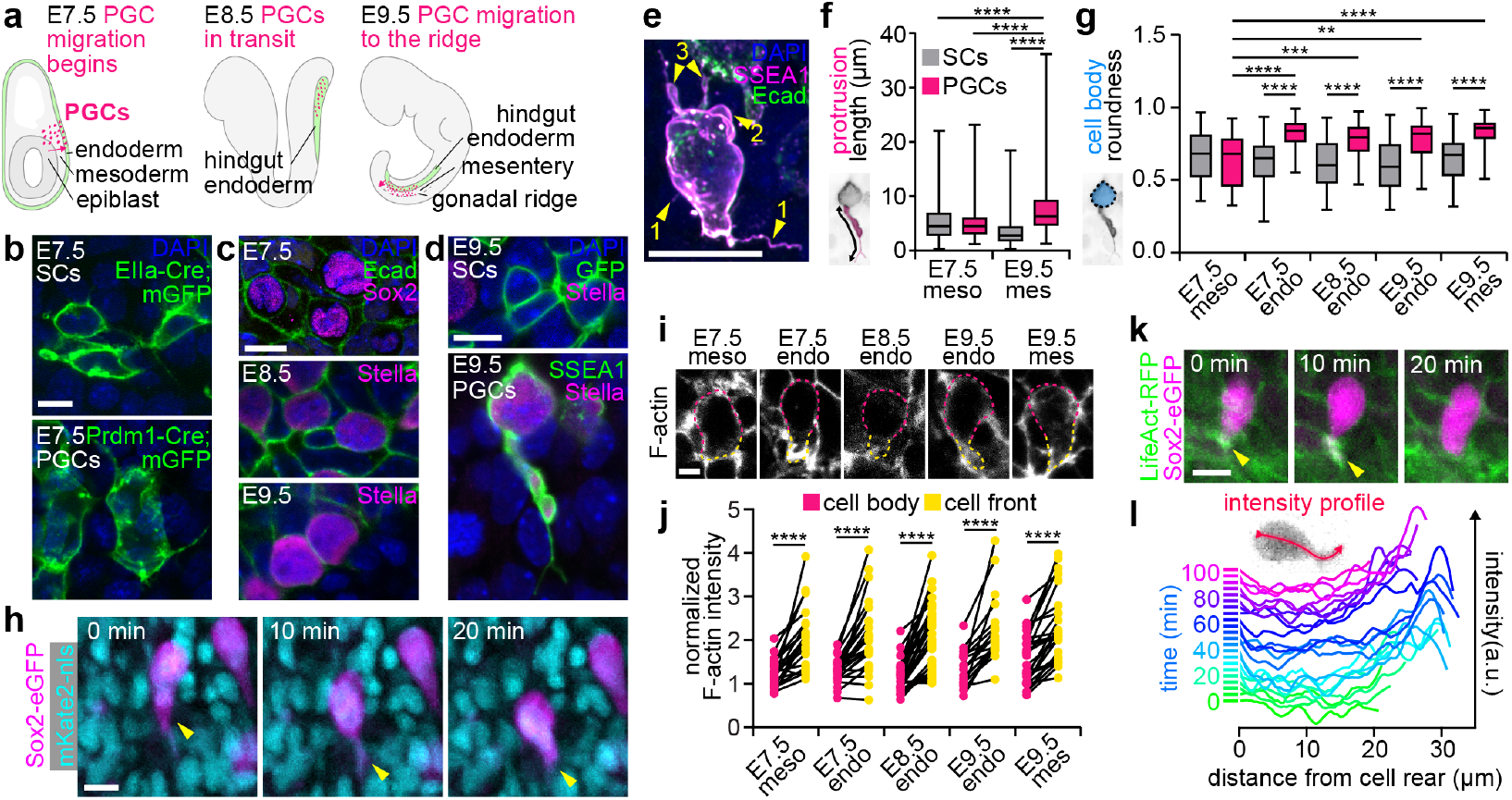
Migrating PGCs extend actin-rich protrusions. **a** PGC migration in the mouse embryo across developmental stages. **b** Mesoderm of E7.5 EIIa-Cre;mTmG (somatic cells [SC]) and Prdm1-Cre;mTmG (PGCs) embryos stained for GFP and DAPI. **c** Endoderm of E7.5-E9.5 wildtype embryos. PGCs are labelled by Sox2 or Stella. **d** Hindgut mesentery of an E9.5 EIIa-Cre;mTmG embryo and E9.5 wildtype embryo with PGCs labelled by Stella and SSEA1. **e** Airyscan image of E9.5 PGC within the hindgut mesentery stained for SSEA1, Ecad, and DAPI, displaying filopodia (arrowheads 1), blebs (2), and cytoplasm-filled protrusions (3). **f-g** Protrusion lengths (f) and roundness of somatic cell and PGC bodies (g), as defined in the schematics. **h** Time lapse of an E9.5 PGC in a Sox2-eGFP;mKate2-nls hindgut undergoing protrusive migration. **i** PGCs in E7.5-E9.5 embryos stained for F-actin (dashed lines indicate cell body and front). **j** F-actin intensity at PGC fronts and bodies normalized to somatic cells. **k** Time lapse of a PGC in a Sox2-eGFP;LifeAct-RFP embryo. Arrowheads indicate actin-rich protrusion. **l** LifeAct-RFP intensity profiles along a migrating PGC over time. **** denotes p < 0.0001, *** p <0.001, ** p <0.01. For all figures, n values and statistical tests are in Extended Data Table 1. Scale bars 10 *µ*m.

PGCs in different species use a variety of migration modes, from mesenchymal to ameboid[11]. Zebrafish PGCs use an ameboid migration mode characterized by a large bleb at the front of the cell with a contractile acto-myosin network around its base[12]. In *Drosophila*, PGCs maintain a rounded shape and migrate using actomyosin contractility-dependent retrograde actin flow along the entire cell length[13]. In the mouse, early electron microscopy studies showed that migrating PGCs make irregular cytoplasmic protrusions or “pseudopods”[14, 15]. Short-term live imaging studies demonstrated that at all stages, PGCs exhibit active migration, characterized by polarized cell shape and a protrusion in the direction of migration[10, 16, 17]. This is more characteristic of a mesenchymal migration mode, which typically involves adhesion to the ECM. In line with this idea, extracellular matrix (ECM) components are found along the PGC migration path in the embryo[18], and loss of *β*1-integrin (*Itgb1*), a cell-ECM adhesion receptor, impairs PGC migration from the hindgut to the gonadal ridge[16]. However, it has yet to be shown definitively which migration mode(s) PGCs use throughout their journey, and whether they use the ECM as a scaffold for migration.

Defects in morphogenesis, for example the development of the hindgut, prevent PGCs from reaching the gonadal ridge[7]. Subtler changes to PGC host tissues also influence PGC migration. Mouse mutants with altered levels of ECM deposition along the PGC migratory path have revealed an inverse relationship between the amount of ECM and PGC migration rate[19, 20]. Increased fibronectin levels impair PGC migration into the endoderm, while decreased collagen I in the hindgut accelerates PGC migration to the gonadal ridge[19, 20]. PGC migration is therefore strongly influenced by the surrounding microenvironment. In zebrafish, PGCs adapt their migration mode in response to environmental changes. Specifically, they exhibit different blebbing behaviour in ectodermal and mesodermal environments in mutants with disrupted germ layer specification[21]. Regardless of migration mode, cells migrating through physically confined environments must also contend with the nucleus, which can hinder migration and is susceptible to damage upon deformation[22, 23]. It is essential that PGCs maintain their genetic integrity as they migrate, and the question of how they adapt to their changing environment while protecting their nucleus remains unanswered.

Here, we performed dynamic and quantitative analyses of PGC migration from E7.5 to E9.5 in the mouse embryo, providing the first comprehensive study of the migratory characteristics of PGCs throughout their journey from their point of origin to the gonads. We found that PGCs migrate using actin-rich protrusions and maintain this migration mode in each tissue they traverse. Throughout their journey, they not only associate with but actively produce their own ECM environment, providing a possible scaffold for their migration. As the tissues around them develop, PGCs experience increasing cell and nuclear deformation due to physical confinement – conditions that cause nuclear rupture and DNA damage in migrating cells in culture[24]. In line with this, we observe higher frequencies of cell rupture and DNA damage at later stages of PGC migration and when confinement is pharmacologically increased. At the same time, PGCs deplete their nuclear lamina, possibly softening the nucleus in order to preserve nuclear integrity during confined migration.

Our study is the first to examine physical mechanisms of PGC migration in the mouse and the role of mechanical signals on mammalian PGCs *in vivo*. We have uncovered a surprising risk to genome integrity in migrating PGCs, with exciting implications for DNA repair and adaptations in nuclear mechanics in PGCs.

## Results

### Protrusive migration in mouse PGCs

Different cell migration modes use a variety of cell shapes and protrusions powered by distinct underlying machinery[11]. To visualize PGC shapes, we labelled PGC membranes at each stage of their migration and in each tissue of residence and compared them to those of neighbouring somatic cells, using strategies relevant to each stage (See Methods; Fig. 1b-d). We observed protrusions from both PGCs and somatic cells in the mesoderm and mesentery at E7.5 and E9.5 (Fig. 1b,d). In particular, a stunning diversity of protrusions were observed by PGCs in the mesentery at E9.5, including fine filopodia, blebs, and cytoplasm filled protrusions (Fig. 1e). Protrusion lengths were similar at E7.5, but by E9.5, PGCs extended longer protrusions than somatic cells (Fig. 1f). Further, the large cytoplasm-filled enriched pro-trusions observed on PGCs at E9.5 were not seen on somatic cells (Fig. 1d). We also observed a clear change in PGC shape compared to somatic cells: by the time PGCs reached the endoderm (E7.5), they took on a more rounded appearance, which persisted as they migrated through the mesentery (E9.5; Fig. 1c-d). Quantification of the cross-sectional area and roundness of PGC and somatic cell bodies (excluding protrusions) showed no significant differences between PGCs and neighbouring somatic cells in the mesoderm at E7.5, but at all other stages, PGCs displayed greater crosssectional areas and increased roundness than their somatic counterparts (Fig. 1g, Supplementary Figure 1b). Live imaging of hindgut explants revealed that PGCs migrate through the mesentery with these long protrusions at the cell front and maintain a mostly rounded cell body (Fig. 1h, Movie 1).

Cells using mesenchymal migration modes tend to have long, actin-rich protrusions, whereas those undergoing bleb-based or ameboid migration tend to remain rounded and accumulate actin at the base of the bleb or towards the rear of the cell[11]. As PGCs exhibit both rounded bodies and long protrusions, we sought to clarify their migration mode by examining F-actin distribution in migrating PGCs. In fixed samples, migrating PGCs often have a clearly identifiable cell front based on their polarized shape (Fig. 1i). Using immunofluorescence, we quantified F-actin intensity along the cortex at the cell front and in the rest of the cell body. At each stage and in each tissue, F-actin was enriched at the cell front relative to the cell body consistent with a migration mode driven by actin-rich protrusions (Fig. 1j, Supplementary Figure 1c). We also carried out live-imaging of hindgut explants expressing Sox2-eGFP, which is expressed in PGCs throughout these stages, and LifeAct-RFP. Consistent with our fixed analyses, we observed LifeAct-RFP enrichment at the PGC front during migration (Fig. 1k, Movie 2). Furthermore, migration followed a characteristic progression: as PGCs extended protrusions and lengthened, LifeAct-RFP intensity increased at the cell front, and as protrusions disappeared and cells shortened, the actin enrichment decreased as well (Fig. 1l). These data show that PGCs migrate using actin-rich protrusions, a migration mode distinct from that used by *Drosophila* and zebrafish PGCs[12, 13, 25]. Moreover, we find that PGCs use actin-rich protrusions throughout their migration, regardless of the surrounding tissue.

Cell migration mode is often influenced by the surround-ing microenvironment, including the ECM. We therefore examined the distribution of ECM in the mesodermal tissues that PGCs migrate through, focusing on ECM components previously studied in the context of PGC migration[18, 26]. At E7.5, collagen IV, fibronectin, and laminin are enriched specifically in the region of the mesoderm containing PGCs at the proximal-most region of the posterior of the embryo (Fig. 2a-b, Supplementary Figure 2a). At E9.5, these ECM components are present throughout the mesentery (Supplementary Figure 2b), consistent with prior reports[18]. At the single-cell level, we found that each ECM component is enriched around PGCs at E7.5 compared to somatic cells, but found at similar levels around both PGCs and somatic cells at E9.5 (Fig. 2c-d, Supplementary Figure 2c-j). Live-imaging with a fluorescently labelled Lamb1-reporter allele (Lamb1-tdTomato) revealed that, at E7.5, PGCs arrive in the posterior mesoderm surrounded by clusters of laminin (Fig. 2e, Supplementary Figure 2k, Movie 3). This close association of PGCs with laminin continues in the later stages of migration, as visualized by two laminin fusion protein reporters, Lamb1-tdTomato and Lamc1-tdTomato, which are both enriched around PGCs in the hindgut endoderm and mesentery (Fig. 2f-h, Supplementary Figure 2k-n). Notably, PGCs are uniquely enriched in Lamc1-tdTomato in the hindgut endoderm, which is absent from the surrounding somatic cells. This enrichment of laminin in PGCs raises the possibility that they are producing their own laminin and thus influencing their own ECM microenvironment. Consistent with this, Airyscan super-resolution microscopy of hindguts at E9.5 revealed the presence of the Lamb1-tdTomato reporter inside PGCs, likely representing ECM that has not yet been secreted (Fig. 2i). Additionally, visualization of mRNA products using *in situ* hybridization chain reaction (HCR) further confirmed that PGCs at all stages of migration and in each resident tissue express Lamb1 (Fig. 2j). The ability to produce their own ECM may enable PGCs to maintain their protrusive migration mode throughout their journey.

**Figure 2:**
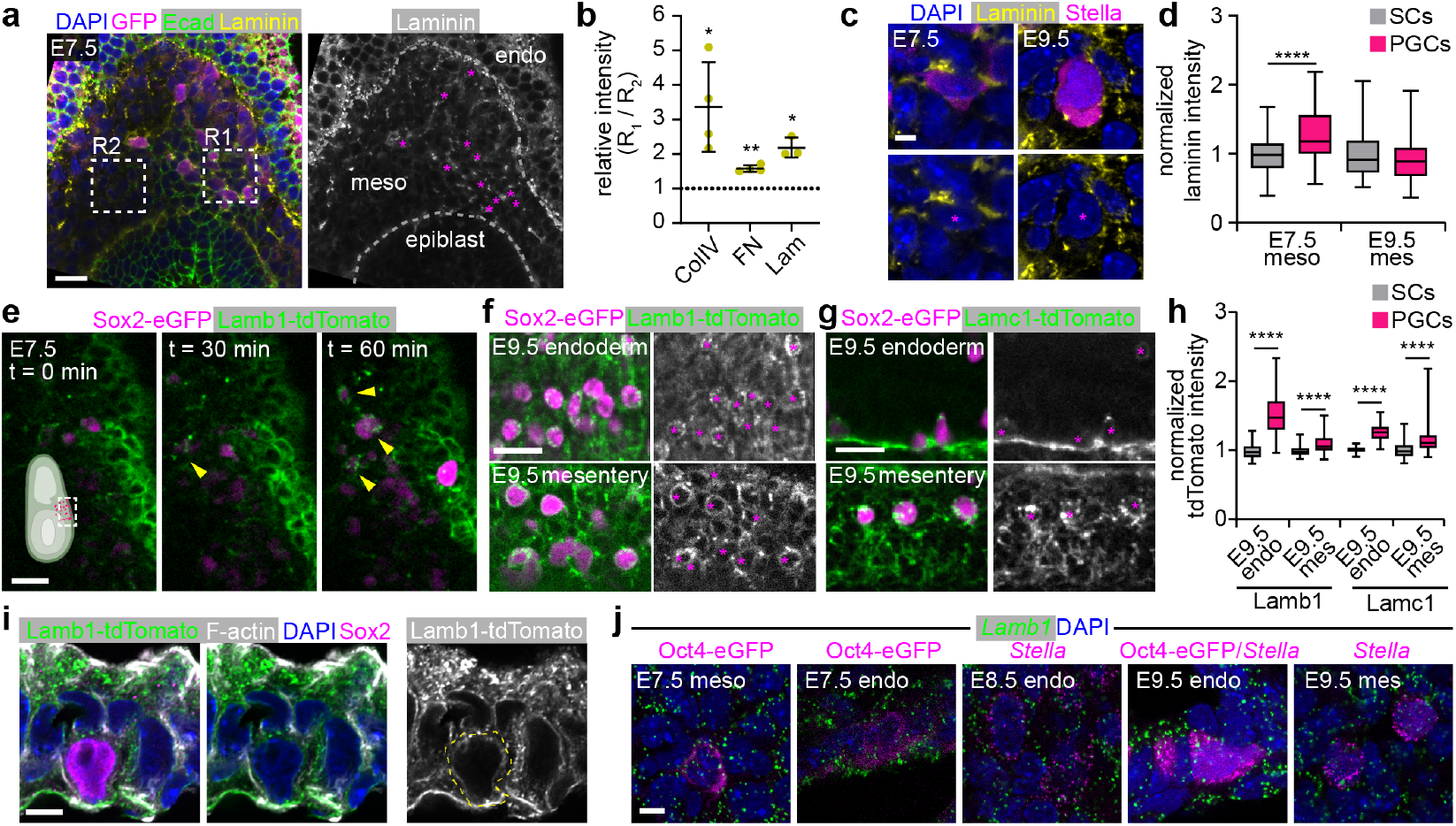
PGCs migrate along ECM-rich paths and produce laminin. **a** Maximum intensity projection of the posterior region of a Sox2-eGFP E7.5 embryo stained for laminin (*Lama1*), GFP, Ecad, and DAPI. Magenta asterisks indicate PGCs. **b** Relative immunofluorescence intensity of ECM components in mesoderm containing PGCs (R1) compared to region without (R2) compared to 1 (i.e., equal intensity). **c** PGCs in E7.5 embryos and E9.5 hindguts stained for laminin (Lama1), Stella, and DAPI. Magenta asterisks indicate PGCs. **d** Immunofluorescence intensity of laminin (*Lama1*) along PGC and somatic cell contours, normalized to somatic cells. **e** Time lapse of an E7.5 embryo expressing Sox2-eGFP and LamB1-tdTomato at the proximal-posterior portion of the embryo as shown in the schematic, including a zoomed-in view of the final timepoint **f-g** E9.5 hindguts from embryos expressing Sox2-eGFP and Lamb1-tdTomato (f) or Lamc1-tdTomato (g) showing PGCs in the hindgut endoderm and mesentery. Asterisks indicate PGCs. **h** Lamb1- and Lamc1-tdTomato intensity around PGCs normalized to somatic cells in each embryo. **i** Airyscan images of sectioned LamB1-tdTomato hindguts. Dashed yellow line indicates PGC outline based on F-actin staining. **j** HCR in situ hybridization of E7.5, E8.5, and E9.5 embryos showing the expression patterns of *Lamb1* and the PGC markers Stella (*Dppa3*) or Oct4-eGFP in the mesoderm (meso) and endoderm (endo). **** denotes p < 0.0001, ** p <0.01, * p <0.05. Scale bars 25 *µ*m in (a), (e), and (f), and 5 *µ*m in (c), (h), and (i).

### Basement membrane formation influences PGC entry to and exit from the endoderm

In addition to migrating through ECM-rich microenvironments, PGCs must also migrate through ECM-enriched barriers at tissue boundaries, suggesting the need to actively remodel the ECM. As PGCs migrate from the mesoderm to the endoderm at E7.5, and then from the endoderm to the mesentery at E9.5, they must traverse the basement membrane of the endoderm. Depending on the structure of the basement membrane, it may act as a physical barrier to PGC migration. We therefore examined the distribution of collagen IV, fibronectin, and laminin in the endodermal basement membrane from E7.5 to E9.5 (Supplementary Figure 3a). Based on each ECM component, the basement membrane of the endoderm is present, but thin and discontinuous, at E7.5 and becomes thicker and more continuous through E9.5 (Supplementary Figure 3b-d). The basement membrane of the endoderm is therefore established during PGC migration and residence in the endoderm.

We hypothesized that at the time of PGC arrival, the basement membrane may not be sufficient to hinder PGC entry into the endoderm at E7.5, but that the more established basement membrane at E8.5 and E9.5 may hinder PGC exit.

We therefore examined PGC shapes upon entry to and exit from the endoderm. At E7.5, before a consistent basement membrane is present, PGCs entering the endoderm show little constriction or deformation (Fig. 3a-b). To visualise these events in live embryos, we used time lapse light-sheet imaging data[17]. Focusing on the proximal posterior region of the embryo around E7.5 allowed us to visualize PGC entry into the outer endoderm layer of the embryo. PGCs were observed to enter the endoderm layer and occasionally exit shortly before re-entering (Fig. 3c). We quantified the distance between PGCs and the surface of the embryo over time and found that many PGCs migrate to and from the endoderm at this stage of development (Fig. 3d). The transient entry of PGCs into the endoderm layer is consistent with our hypothesis that the nascent basement membrane at E7.5 does not act as a significant barrier to PGC movement.

**Figure 3:**
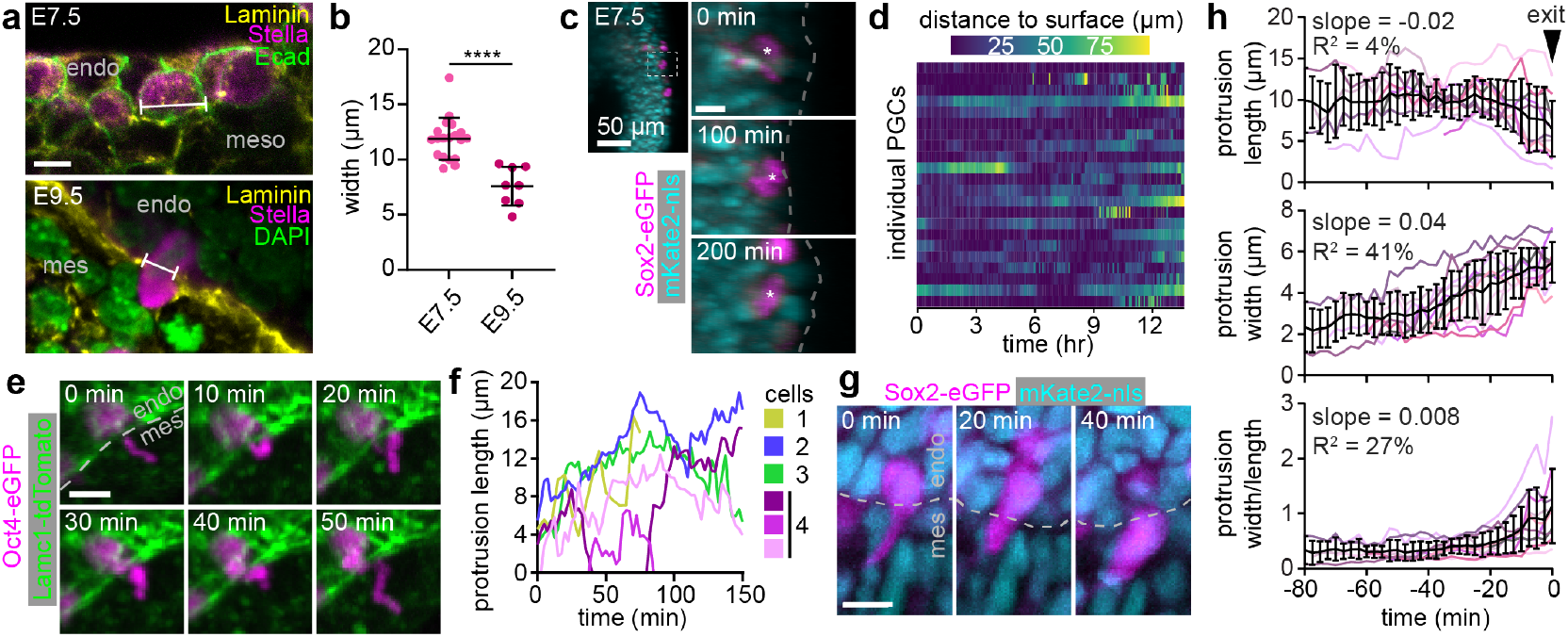
Basement membrane formation influences PGC shapes and dynamics upon entry to and exit from the endoderm. **a** Posterior region of an E7.5 embryo and E9.5 hindgut stained for Laminin, Stella, and either Ecad or DAPI to show PGC morphology during entry to (E7.5) and exit from (E9.5) the endoderm. Location of width measurement indicated. **b** Width of PGCs measured at the level of the basement membrane as they cross into or out of the endoderm at E7.5 and E9.5, respectively. **c** PGC transiently entering the endoderm in an E7.5 embryo expressing Sox2-eGFP and mKate2-nls. Asterisk indicates PGC and dashed line indicates embryo surface **d** Distance of individual PGCs to the surface of the embryo over time. **e** PGC protrusion dynamics in a E9.5 hindgut explant expressing OCT4-eGFP and Lamc1-tdTomato. **f** Sample traces of protrusion length over time. **g** PGC exiting the hindgut endoderm in a Sox2-eGFP;mKate2-nls hindgut explant. **h** Protrusion length, width, and ratio of width to length during PGC exit from endoderm. Time courses are aligned based on when the PGC nucleus is no longer in the endoderm. Mean and standard deviation (black) are shown along with individual samples. **** denotes p < 0.0001. Scale bars 10 *µ*m unless otherwise indicated.

In contrast to the rounded shapes of PGCs entering the endoderm at E7.5, PGCs migrating across the thicker basement membrane at E9.5 exhibit more deformation (Fig. 3a-b), suggesting they must squeeze through the basement membrane to exit. Further, unlike PGCs at E7.5, we did not observe any PGCs re-entering the endoderm after they had crossed the basement membrane into the mesentery at E9.5. The establishment of the endoderm basement membrane may therefore act as a barrier to PGC exit during hindgut involution that PGCs must then overcome at E9.5 to continue their migration. To understand how PGCs accomplish this exit event, we examined fixed hindguts and performed live-imaging. At E9.5, we observed PGCs whose cell bodies were still in the endoderm projecting long, thin protrusions into the mesen-tery (Fig. 3e). Using time lapse imaging of hindgut explants, we found that these protrusions were highly dynamic, could rapidly extend and retract, and that PGCs could extend multiple protrusions simultaneously (Fig. 3e-f, Movie 4). These protrusions might play a role in PGC exit from the endoderm, either in exploring for a suitable path to migrate along, or by opening a hole in the basement membrane large enough for the cell body to squeeze through. In line with this idea, tracking exiting PGCs in hindgut explants revealed that these protrusions precede PGC exit from the hindgut endoderm (Fig. 3g, Movie 5). Protrusions followed a similar progression: a long, thin, and long-lived protrusion eventually widened immediately prior to PGC exit from the endoderm (Fig. 3h).

### The developing embryo imposes increasing confinement on migrating PGCs

The effects of basement membrane development on PGC migration dynamics suggests that the properties of the tissue microenvironment may influence the speed or success of PGC migration to the gonadal ridges. We therefore examined tissue-level physical properties around PGCs at each stage of development. We compared the sizes of intercellular spaces with those of PGCs by incubating embryos and hindgut explants with 70 kDa dextran. We found that intercellular spaces in the tissues surrounding PGCs were significantly smaller than the PGCs themselves, and that they decrease as migration proceeds from E7.5 to E9.5 (Fig. 4a, Supplementary Figure 4a-b). To estimate the elastic moduli of tissues that PGCs migrate through (as a measure of tis-sue stiffness), we performed atomic force microscopy (AFM) on embryonic tissues from E7.5 to E9.5. Tissue stiffness was first low in the mesoderm and endoderm at E7.5, and then increased gradually in the hindgut endoderm through E9.5 (Fig. 4b, Supplementary Figure 4c-d). At E9.5, when PGCs exit the endoderm, the hindgut mesentery was less stiff than the endoderm, but stiffer than the mesoderm and endoderm at E7.5 and the endoderm at E8.5 (Fig. 4b, Supplementary Figure 4c-d). As cells tend to migrate along stiffness gradients in culture and *in vivo*[27, 28], we examined the path through the mesentery to the ridge but found that tissue stiffness remained constant (Fig. 4b, Supplementary Figure 4c-d). Given the decrease in intercellular space and the increase in tissue stiffness from E7.5 to E9.5, we conclude that the tissues around migrating PGCs impose increasing confinement over the course of migration.

**Figure 4:**
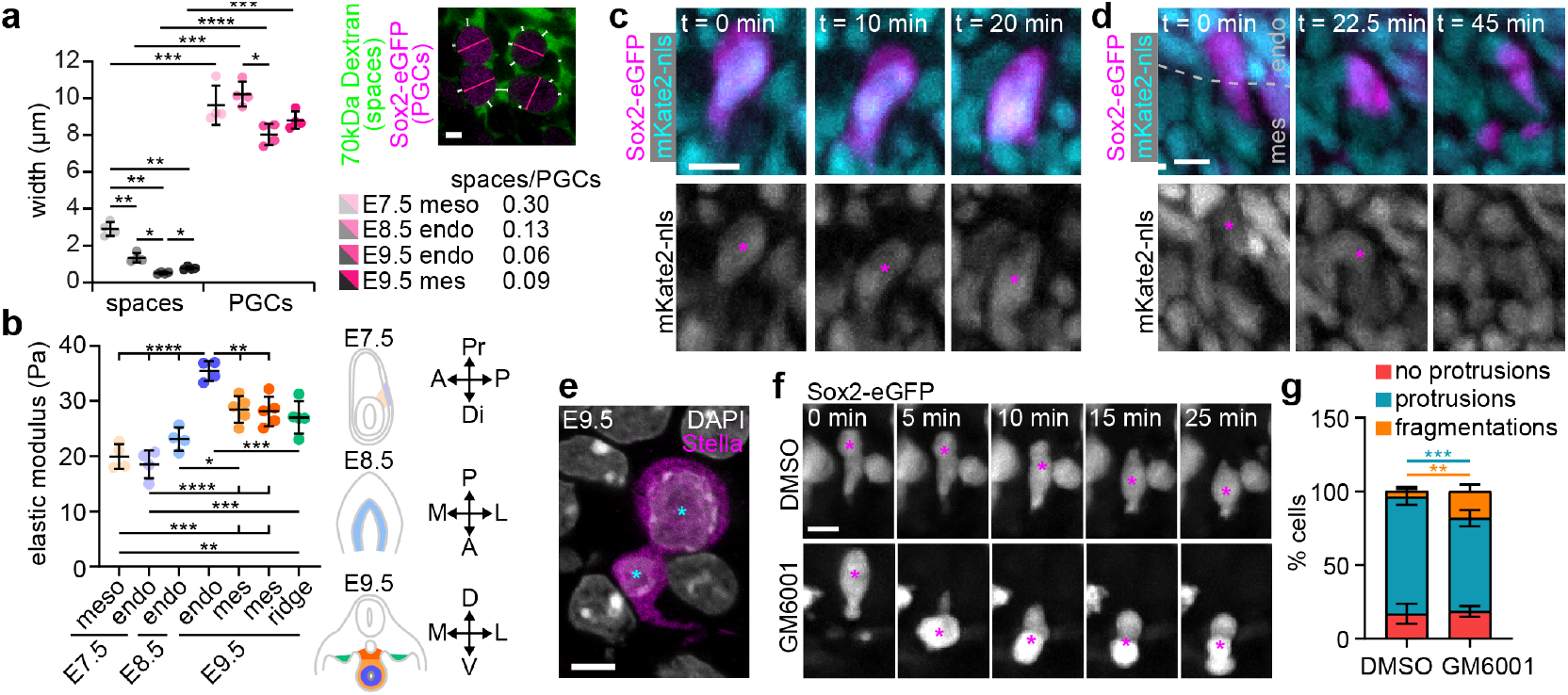
Confined migration leads to cell and nuclear rupture in migrating PGCs. **a** Average widths of intercellular spaces (measured using fluorescent Dextran as indicated in image) and PGCs. **b** AFM measurements of the average elastic modulus of tissues around migrating PGCs from E7.5 to E9.5, as indicated by the schematics (A: anterior, P: posterior, Pr: proximal, Di: distal, M/L:medial-lateral, D: dorsal, V: ventral). **c** Time lapse of an E9.5 Sox2-eGFP;mKate2-nls hindgut explant showing nuclear and cell body deformation in a PGC migrating through the mesentery. **d** Time lapse of an E9.5 Sox2-eGFP;mKate2-nls hindgut explant showing a PGC exiting the hindgut endoderm and immediately fragmenting. **e** Fragmented nucleus of a PGC in the hindgut mesentery of an E9.5 embryo stained for Stella and DAPI. Cyan asterisks indicate the two nuclear fragments. **f** Time lapse of PGCs in the mesentery of an E9.5 Sox2-eGFP-expressing hindgut explant treated with GM6001 and DMSO control. **g** Quantification of cell behaviours in GM6001-treated and control explants. **** denotes p < 0.0001, *** p <0.001, ** p <0.01, * p < 0.05. Scale bars 5 *µ*m in (a) and (e), 10 *µ*m in (c), (d), and (f).

As a result of this increasing confinement and the restrictions imposed by the basement membrane, PGCs and their nuclei must deform significantly in order to exit the hindgut endoderm and move through the mesentery (Fig. 4c, Movie 1). *In vitro* cell culture work has demonstrated that cancer cells migrating through narrow microfluidic channels experience nuclear rupture and DNA damage[24]. We therefore asked whether a similar phenomenon occurs in migrating PGCs. We were able to identify clear cases of cell rupture and death. In some cases, PGC deformation upon exit from the endoderm is followed by immediate fragmentation of the PGC (Fig. 4d, Movie 6). We also observed rare cases of PGCs with severely disrupted nuclei, including a dramatic example of a PGC whose nucleus had been split in two (Fig. 4e). To de-termine whether these cell fragmentation events were linked to confinement, we tuned the extent of confinement by inhibiting matrix metalloproteinase (MMP) activity. As migratory PGCs express a repertoire of MMPs[29], we hypothesized that treatment with the MMP inhibitor GM6001 would prevent ECM remodelling and make it more difficult for PGCs to migrate through the mesentery. In GM6001-treated explants, intercellular spaces are significantly narrowed, leading to increased confinement (Supplementary Figure 4e-g). While GM6001 treatment did not affect the movement or protrusive activity of PGCs, we observed a significant increase in the number of fragmentation events (Fig. 4f-g). In particular, we observed PGCs deforming rapidly and dramatically while squeezing through the tissue, followed by the sudden fragmentation of the cell (Fig. 4f, Movie 7). These data suggest that confinement in the mesentery can cause mechanically-induced cell rupture in PGCs, and that ECM remodelling is important for PGCs to migrate through the mesentery without incurring damage.

Given that PGCs experience significant deformations and even cell rupture under confinement, we next asked whether PGCs incur DNA damage as they migrate. We used three different markers for DNA damage (*γ*H2AX, RPA32, and 53BP1) and compared their levels in PGCs to those in the surrounding somatic cells in the mesoderm, endoderm, and mesentery at E7.5, E8.5, and E9.5. At E7.5, PGCs and somatic cells of the mesoderm and endoderm did not display any accumulation of damage (Fig. 5a-b, Supplementary Figure 5a-g). However, starting at E8.5 and increasing as PGCs travel from the endoderm to the mesentery at E9.5, PGCs accumulated DNA damage at levels greater than the surrounding somatic cells (Fig. 5a-b, Supplementary Figure 5a-g). These data show that DNA damage indeed becomes more prevalent in migrating PGCs as they are subjected to increased confinement from the developing tissues and basement membranes of the embryo. To further confirm this, we treated hindgut explants with GM6001 to prevent matrix remodelling and increase confinement, and found increased incidence of DNA damage in PGCs, but not in neighbouring somatic cells (Fig. 5c-d, Supplementary Figure 5h-k). With this evidence we conclude that PGCs migrating *in vivo* experience confinementinduced DNA damage.

**Figure 5:**
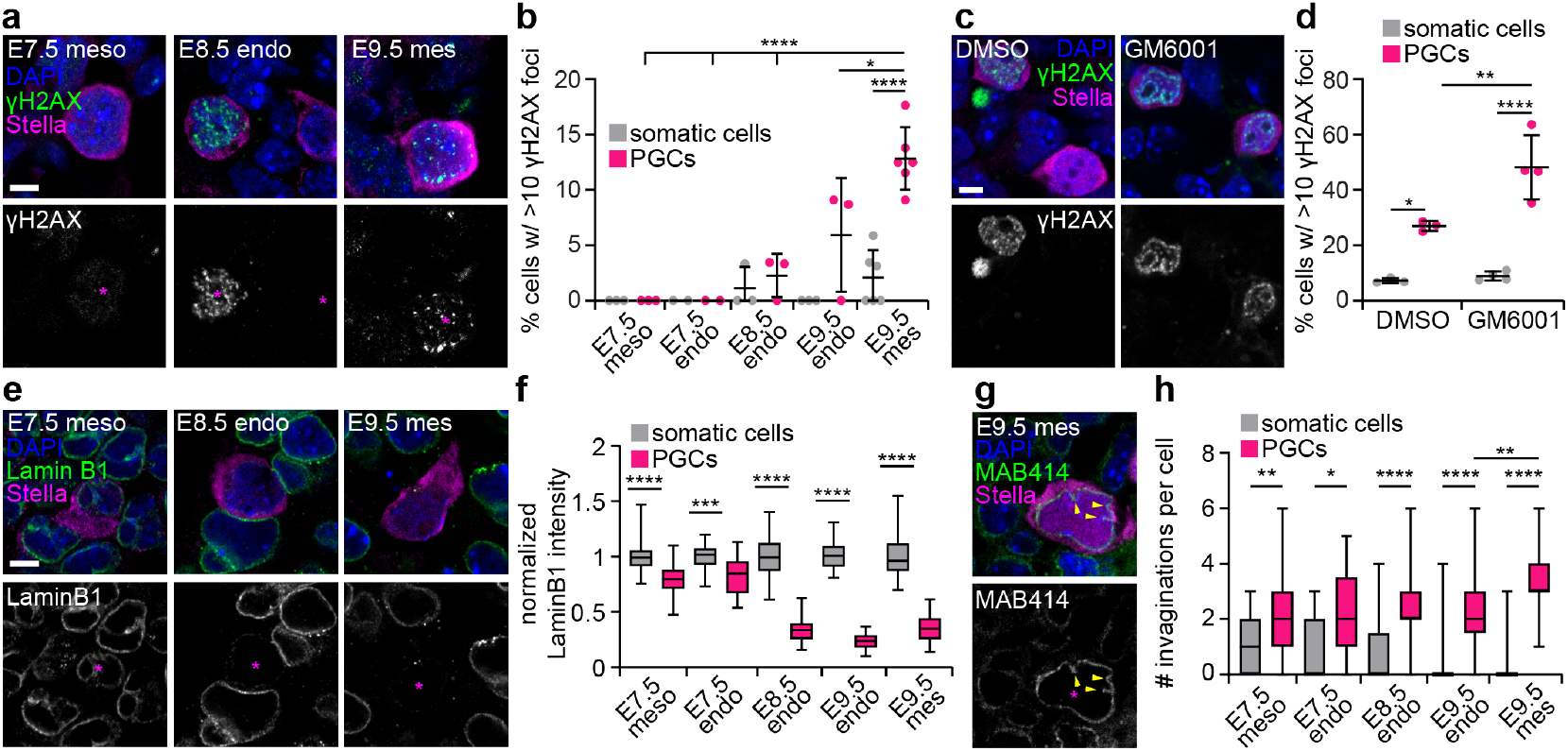
PGCs exhibit increasing DNA damage and remodel their nuclear lamina throughout migration. **a** Sections of E7.5, E8.5, and E9.5 embryos stained for *γ*H2AX, Stella and DAPI. Asterisks indicate PGCs. **b** Percentage of cells with >10 *γ*H2AX foci in PGCs and in surrounding somatic cells. **c** Sections of E9.5 hindguts cultured with DMSO or GM6001 and stained for *γ*H2AX, Stella and DAPI. Asterisks indicate PGCs. **d** Percentage of cells with >10 *γ*H2AX foci in PGCs and in surrounding somatic cells in GM6001-treated explants and DMSO controls. **e** Sections of E7.5, E8.5, and E9.5 embryos stained for Lamin B1, Stella and DAPI. **f** Immunofluorescence intensity of Lamin B1 around the nuclei of PGCs and somatic cells normalized to somatic cells. **g** Sections of E9.5 embryos stained for MAB414, Stella and DAPI. Yellow arrowheads indicate invaginations of the nuclear envelope. **h** Number of invaginations of the nuclear envelope in PGCs and in surrounding somatic cells. **** denotes p < 0.0001, *** p <0.001, ** p <0.01, * p < 0.05. Scale bars 5 *µ*m.

It is surprising that cells as essential as PGCs must undertake a migration that can lead to DNA damage and cell death. We therefore wondered whether PGCs had any compensatory mechanisms that could help them cope with these stresses. Softening of the nuclear envelope, for example, might help PGCs squeeze their nuclei through narrow spaces without inducing rupture. Nuclear lamins are known to affect stiffness and deformability of the nucleus[30]. We therefore examined nuclear lamins in PGCs over the course of their migration. Lamin A/C is absent from all cells of the embryo until approximately E9.5[31], and we confirmed that this was also true for PGCs (Supplementary Figure 6a-b). Immunofluorescence analysis of Lamin B1 at the nuclear periphery showed that PGCs have slightly lower Lamin B1 levels than surrounding somatic cells at E7.5 (Fig. 5e-f, Supplementary Figure 6c-d). However, from E8.5 through E9.5, in both the endoderm and mesentery, Lamin B1 levels are dramatically reduced in PGCs compared to neighbouring somatic cells (Fig. 5e-f, Supplementary Figure 6c-d). Throughout these stages, PGCs also have more Lamin B2 at the nuclear periphery than somatic cells, but Lamin B2 appears at low levels and with a punctate distribution along the nuclear envelope (Supplementary Figure 6e-g). These changes in the nuclear lamina could give rise to altered mechanical properties of the nucleus. Consistent with this, PGCs have more deformed nuclear membranes than somatic cells throughout their migration, as evidenced by an increase in the number of nuclear envelope invaginations (Fig. 5g-h, Supplementary Figure 6h-i).

## Discussion

Here, we carried out *in vivo* analyses of the physical mechanisms of mouse PGC migration and investigated how PGCs are influenced by the changing mechanical environment of the embryo. We found that regardless of the surrounding tissue, PGCs maintain the use of actin-rich protrusions to migrate. Starting at the earliest stages and continuing throughout migration, PGCs retain close associations with ECM and express laminin. However, as the embryo develops, certain aspects of PGC behaviour adjust. The developing basement membrane acts as a barrier to PGC exit from the hindgut endoderm, forcing PGCs to extend and then expand protrusions through the basement membrane in order to migrate into the mesentery. We demonstrate that development of the hindgut endoderm and the mesentery leads to increasing confinement and mechanical stiffness, forcing migrating PGCs and their nuclei to undergo significant deformations. Cell rupture events occur among PGCs migrating through these confined spaces, and these events become more frequent when confinement is pharmacologically increased. In the later stages when confinement is highest, PGCs display higher amounts of DNA damage than in the early stages of migration or than their somatic neighbours. PGC-specific DNA damage is further amplified when confinement is pharmacologically increased. Concurrent with the onset of migration, PGCs deplete Lamin B1 and develop wrinkled nuclear envelopes, possibly indicating a softer nucleus which could ease the effects of confined migration and prevent mechanical damage to the nucleus. Overall, we demonstrate that migrating PGCs in the mouse employ a migration mode that is maintained irrespective of their surroundings, but as the embryo develops, increasing stiffness and confinement pose potential risks to PGCs and their nuclei.

The combination of PGCs forming actin-rich protrusions at the cell front, moving along ECM-rich routes, and reliance on MMPs are all features of a mesenchymal migration mode[11]. Our observations suggest that PGCs in the mouse migrate using a distinct strategy to that employed by PGCs in *Drosophila* or zebrafish[4, 12, 13, 25], highlighting the importance of studying this process in a mammalian model. Mesenchymal migration also usually involves strong and specific adhesion to a substrate. Since PGCs in the mouse pass through different tissue types, with different available ECM and cell-cell adhesions, PGCs may adapt specific adhesions to their host tissue. In mesodermal tissues, the ECM provides a possible scaffold for migration anchored by integrin-mediated adhesions[18, 26], while Ecad-mediated adhesions could play a similar role in the endoderm, as has been shown for PGCs in zebrafish[25]. Alternatively, PGCs may use non-specific adhesion, which would enable them to maintain the same migration machinery regardless of the surrounding tissue[32]. To fully elucidate the mechanisms of PGC migration, future work will need to determine the requirement (or lack thereof) of different adhesion complexes in each tissue PGCs encounter.

In cells migrating under confinement, squeezing the nucleus through tight spaces often acts as a rate limiting step[33]. The effect of the nucleus in these contexts depends on nuclear mechanical properties – for example, depletion of Lamin A allows cells in culture to migrate through narrow channels much faster[34]. Here, we show that PGCs, which lack Lamin A/C, deplete Lamin B1 during migration stages and have minimal punctate Lamin B2 along the nuclear envelope. Depletion of the nuclear lamina likely results in a more flexible nucleus, in line with our observations of PGC nuclear envelope shape and invaginations. These changes to the nucleus may allow PGCs to migrate under confinement more quickly and decrease the chance of incurring DNA damage. PGC migration in the mouse may therefore be an exciting model for investigating how the nucleus adapts during migration *in vivo*. Beyond the nuclear lamina, nuclear mechanical properties are also affected by chromatin organization[35, 36]. Throughout migratory stages, PGCs undergo significant epigenetic remodelling and changes in chromatin organization[37, 38]. Future work should determine how these changes to chromatin, along with depletion of lamins, affects the mechanical properties of the PGC nucleus, the rate of migration, and the incidence of DNA damage. Our work suggests that mammalian PGCs adapt in surprising ways to the developing tissues around them: they maintain their migration strategy, but exhibit changes at the level of the nucleus that may facilitate successful, damage-free migration to the future gonads.

The later stages of PGC migration are accompanied by increased incidence of DNA damage specifically in PGCs, and pharmacologically increased confinement amplifies this damage. We propose that this damage is incurred due to the increasingly stiff and confined environment around PGCs. Nuclear deformation associated with confined migration and nuclear rupture, as well as nuclear compression leading to replicative stress, are both well documented causes of DNA damage in culture[24, 39]. However, to our knowledge, this study provides the first possible case of this phenomenon in a mammalian embryo. Given the importance of genome integrity to the germline, our findings raise important questions about whether and how DNA damage in migrating PGCs is repaired or if damaged PGCs are later excluded from the germline through apoptosis. Mutant mice lacking components of certain DNA damage repair pathways show striking PGC-specific defects in the embryo[39, 40]. While somatic lineages are unaffected, PGCs in the immediate postmigratory stages accumulate DNA damage and are eliminated, leading to fewer PGCs as early as E9.5 and reduced fertility[39, 40]. These findings suggest PGCs are more sensitive to loss of DNA damage repair machinery than other cells in the embryo, and raises the question of how this damage occurs. Our findings suggest that confinement may be a source of DNA damage in migrating PGCs – damage which is normally repaired in non-mutant animals. Whether this repair results in mutations that persist in the germline is a fascinating area for future work.

## Acknowledgments

K.G. and K.M. are supported by the Medical Research Council as part of UK Research and Innovation (MCUP1201/23). This work was supported by the Medical Research Council, as part of United Kingdom Research and Innovation (also known as UK Research and Innovation) [MCUP1201/23]. For the purpose of open access, the MRC Laboratory of Molecular Biology has applied a CC BY public copyright licence to any Author Accepted Manuscript version arising. S. J. A is supported by the German Research Foundation (DFG) through the Heisenberg Program (AR 732/4-1), project grant (AR 732/2-1), project A03 of CRC 850 (project ID 89986987), project A08 of CRC 992 (project ID 192904750) and Germany’s Excellence Strategy (CIBSS – EXC-2189 – Project ID 390939984).

## Methods

### Experimental model and subject details

#### Mice

All animal experiments performed in this study were approved by the Medical Research Council’s Laboratory of Molecular Biology animal welfare and ethical review body and conform to the UK Home Office Animal (Scientific Procedures) Act 1986 (License noPP5976836). The mice used in this study were Blimp1-mGFP1[1], wild-type Hsd:ICR (CD-1), EIIa-Cre (JAX #003724), Ecad-eGFP[41], Oct4-eGFP[16] (JAX #004654), Lamb1-tdTomato (see below), Lamc1-tdTomato (see below), LifeAct-RFP[42], R26-CAG-nuc-3xmKate2-nls (RIKEN)[43], ROSA26 mT/mG (JAX #007576), Prdm1-Cre (JAX #008827), and Sox2-eGFP (JAX #017592). Timed pregnant females were obtained from natural matings, with the presence of a copulatory plug denoted as E0.5.

#### Generation of Lamb1tdTom and Lamc1tdTom reporter strains

A detailed description of the generation and characterization of basement membranes reporter strains in mouse will be reported elsewhere (please contact SJA for more detailed information or requests for the strains). In brief, targeting vectors contained 400-500 bp homology regions on both sides surrounding the STOP-codons contained in the last exons of the Lamb1 (exon 33) and Lamc1 (exon 28) genes. Homologous recombination generates C-terminal fusions of the laminin chains to a 30 aa linker and the tandem dimer tomato fluorescent protein (tdTomato), followed by the SV40 polyadenylation signal. For positive selection targeting vectors additionally contained a PGK.neo positive selection cassette. To increase targeting efficiency, pairs of TALENs were designed to generate 100-200 bp deletions in the 3’UTR of laminin chains. Linearized targeting vector and TALEN pairs were co-electroporated in CCE mouse embryonic stem cells and G418 drug-resistant embryonic stem cell colonies were screened by PCR across the recombination junctions on both 5’ and 3’ sites, and expression of the reporters controlled by fluorescent observation. Correctly targeted embryonic stem cell clones were used for the generation of chimeras. The PGK.neo cassette was removed after successful germline transmission by crossing to the PGK.Cre transgenic line. Primers for genotyping Lamb1-tdTomato were AGCAGCCCTATATCCTCCCT (rev), AGGTTCGCTCCCTCCTTAAG (fw wildtype), TCACTGCATTCTAGTTGTGGT (fw flox). Primers for genotyping Lamc1-tdTomato were AGGGTGCTGACAGAAGTGGA (rev) and CCAGAGGCCACTTGTGTAGC (fw).

### Method details

#### Immunofluorescence staining and imaging

At the desired embryonic stage, timed-pregnant females were euthanized by cervical dislocation and embryos dissected in PBS containing 5% FBS (Corning 35-016-CV). Embryos were then fixed overnight in 4% PFA in PBS. Hindgut dissections were carried out after fixation. For generating tissue sections, E7.5 embryos were frozen in OCT (VWR 361603E) directly, and older embryos were first taken through a sucrose gradient (20%, 30%, 30% sucrose:OCT 1:1) before embedding and freezing. Sections were made on a Leica CM1950 cryostat. Slides were permeabilized in PBS with 0.1% Triton-X, washed with PBS, then incubated in blocking solution (PBS with 0.1% Triton-X and 10% FBS).

Slides were then incubated with primary antibodies in blocking solution overnight at 4°C, washed in PBS, and incubated with secondary antibodies and DAPI (1:1000) diluted in blocking solution for 3 hours at room temperature. Prior to imaging, ProLong Gold antifade reagent (Invitrogen P36930) and a coverslip were added to slides.

Wholemount samples were permeabilized in PBS with 0.5% Triton-X for one hour at room temperature, then incubated in blocking solution overnight at 4°C. Samples were then incubated overnight at 4°C with primary antibodies diluted in the same blocking solution, washed in PBS with 0.1% Triton-X for an hour at room temperature, then incubated overnight at 4°C with secondary antibodies and DAPI (1:1000) diluted in blocking solution. Samples were then washed in PBS with 0.1% Triton-X and transferred to PBS for mounting and imaging.

The primary antibodies and concentrations used are as listed: 53BP1 (rabbit, 1:200, Novus Biologicals NB100-304), cleaved caspase 3 (rabbit, 1:200, CST 9664), collagen IV (goat, 1:200, Millipore AB769), E-cadherin (rat, 1:400, ThermoFisher Scientific 13-1900), *γ*H2AX (rabbit, 1:400, CST 2577), GFP (chicken, 1:400, Abcam ab1397), fibronectin (rabbit, 1:200, Rockland 600-401-117-0.1), integrin *β*1 (rat, 1:200, Merck MAB1997), lamin A/C (chicken, 1:1000, Novus Biologicals NBP2-25152), Lamin B1 (rabbit, 1:500, Abcam ab16048), Lamin B2 (rabbit, 1:200, Abcam ab 151735), laminin (rabbit, 1:200, Sigma-Aldrich L9393), MAB414 (mouse, 1:200, Abcam ab24609), RPA32 (rat, 1:100, CST 2208), Sox2 (rabbit, 1;200, Abcam ab97959), SSEA1 (mouse, 1:200, Novus Biologicals MC-480), Stella (goat, 1:100, R&D Systems AF2566), td-Tomato (goat, 1:400, Antibodies Online ABIN6254170). Secondary antibodies were AlexaFluor conjugated and raised in donkey (1:400, Invitrogen A21208, A21206, A21202, A11055, A11058, A21209, A32754, A21203, A48272, A31573, A32787, A78948). For F-actin staining, Phalloidin (1:400, Invitrogen A22287) was added along with secondary antibodies. DAPI (1:1000, Sigma-Aldrich MBD0015) was also added along with secondary antibodies.

E7.5 embryos were mounted in ProLong Gold antifade reagent (Invitrogen P36930) sandwiched between two glass coverslips separated by a thin silicone barrier to prevent sample deformation. In some cases, they were first sliced open with a fine surgical blade (Fine Science Tools 10316-14) to enable imaging deeper into the embryo. E8.5 embryos and E9.5 hindguts were transferred to glass-bottom dishes for imaging. Samples were imaged on Zeiss 710 or Zeiss 780 inverted laser scanning confocal microscopes with 10×, 20×, 40× water, or 63× oil objectives.

For wholemount AiryScan imaging, samples were cleared by first dehydrating through an isopropanol series (25%, 50%, 75%, and 100%), then transferring to 50% isopropanol in a 1:2 mixture of benzyl alcohol and benzyl benzoate (BABB), then into 100% BABB. Samples were imaged on a Zeiss 880 or Zeiss 900 inverted laser scanning confocal microscope with AiryScan with a 63× oil objective.

Time lapse movies (Movies 1, 2, 4, 5, 6 and 7) and still images captured from these datasets (Figures 1h, k, 3e,g, 4c,d, and f) were denoised using ndsafir[44] with the following parameters applied equally to the entire time series: patch size (9×9), mode (2D+time), smoothing (0.5), iterations (4), sensor gain (2), sensor offset (100), readout noise (2), remix (0.7).

#### HCR *in situ* hybridization

At E7.5, E8.5, or E9.5, timed-pregnant females were euthanized by cervical dislocation and embryos dissected in ice cold 4% PFA. Embryos were then fixed overnight in 4% PFA in PBS. For E8.5 embryos, the posterior-most part of the embryo was isolated by cutting with a fine surgical blade. For E9.5 embryos, cross-sectional segments through the hindgut region were made using a fine surgical blade. Samples were then processed following the standard HCR protocol provided by Molecular Instruments. The probes used were Dppa3-B4 and Lamb1-B2, with B4-488 and B2-647 amplifiers. Prior to imaging, samples were mounted in ProLong Gold antifade reagent (Invitrogen P36930) sandwiched between two glass slides separated by a thin silicone barrier to prevent sample deformation.

#### Embryo and explant culture

At E7.5, E8.5, or E9.5, timed-pregnant females were euthanized by cervical dislocation and embryos dissected in Fluorobrite DMEM (Gibco A18967-01) containing 5% FBS. Embryos or hindguts were cultured in 50% rat serum in Fluorobrite DMEM supplemented with Pen-Strep, GlutaMax (Gibco 35050-061), and MEM non-essential amino acids (Gibco 11140-035). Samples were cultured in glass-bottom 8-well plates (Ibidi IB-80807) and immobilized either by lowering the level of the media such that the sample was trapped in the meniscus at the centre of the well or by adhering to nitrocellulose membranes (Millipore AABG01300). For E7.5 embryos, the ectoplacental cone was adhered to the membrane and for E9.5 hindgut explants, extra tissue on either end of the hindgut was used. To inhibit MMP activity, 10 *µ*M GM6001 (TargetMol T2743) dissolved in DMSO was added to the culture medium, with controls receiving the same volume of DMSO for a final 0.2% DMSO in media. For culture experiments examining DNA damage, hindguts were fixed after 4 hours of incubation with either DMSO or 10 *µ*M GM6001.To visualize intercellular spaces, 70 kDa Dextran (Invitrogen D1830) was added to the culture medium at 1:40. To measure intercellular spaces in GM6001 cultures, hindguts were incubated with DMSO or 10 *µ*M GM6001 and 70kDa Dextran for an hour prior to imaging. For time lapses, samples were imaged on a Nikon CSU-W1 spinning disk confocal microscope equipped with a 25× silicone immersion objective and stage heater. For measuring intercellular spaces, samples were imaged on a Zeiss 780 inverted laser scanning confocal microscope with a 40× water objective.

#### AFM of tissue sections

At E7.5, E8.5, or E9.5, timed-pregnant females were euthanized by cervical dislocation and embryos dissected in Fluorobrite DMEM (Gibco A18967-01) containing 5% FBS. Embryos were then immediately embedded in OCT and frozen on dry ice. 20 *µ*m-thick sections were made on a Leica CM1950 cryostat. Sectioned samples were then kept submerged in PBS for the remainder of the experiment, and all measurements were made within an hour after sectioning. AFM measurements were made using an Asylum MFP3D atomic force microscope operated in contact mode. We used a spherical silicon dioxide tip with a diameter of 6.62 *µ*m and a force constant of 0.08 N/m (Apex Probes CP-PNPL-SiO-C-5) which was calibrated using thermal oscillation in air prior to the experiment. Force curves were obtained using a maximum force setpoint at up to 6 different locations within the regions of interest (E7.5 mesoderm, E7.5 endoderm, E8.5 endoderm, E9.5 mesentery – close and far from the endoderm, and E9.5 genital ridge).

### Quantification and statistical analyses

#### Image analysis

Cell shapes and protrusions were quantified manually in Fiji (Fig. 1f-g, Extended Data Fig. 1a). Specifically, cross-sectional area and roundness were quantified at the confocal plane corresponding to the middle of each cell, and the contours traced excluded any protrusions as shown in Fig. 1f-g. All PGCs in each image were measured, and an equal number of randomly selected nearby somatic cells were also quantified for comparison. These analyses were made using a variety of labelling strategies for cell membranes and protrusions (where possible). For somatic cells at E7.5 and E9.5, we used the mosaic activity of EIIa-Cre in mTmG mice and looked for embryos with sparse enough labelling to distinguish protrusions. For PGCs at E7.5, we used Prdm1-Cre with mTmG and measured only cells within the mesoderm since Prdm1-Cre is also active in the endoderm at these stages. This precluded using Prdm1-Cre for visualization of PGC shapes in the endoderm, so for both PGCs and somatic cells in the endoderm at E7.5, E8.5, and E9.5, we relied on Ecad immunostaining to measure cell shapes. We do not currently have a method for visualizing fine protrusions at these stages. Finally, for PGCs at E9.5, we used SSEA1 immunostaining.

To measure changes in F-actin accumulation and distribution in fixed samples (Fig. 1j, Extended Data Fig. 1c), we measured fluorescence intensity of F-actin (based on phalloidin staining). For PGCs, we measured intensity in a 20 pixel-thick line traced just inside the cell membrane, either at the cell front or in the cell body, as shown in Fig. 1i, in PGCs where a distinct cell front could be seen. For somatic cells, we did not distinguish between a cell front and cell body as they are not migrating directionally, and simply performed the same measurement within the entire cell contour. To control for intensity differences between samples and within samples at different depths of the confocal z-stack, we normalized the intensity measurements for PGCs to that of neighbouring somatic cells. For analysis of LifeAct-RFP (Fig. 1l), we manually traced along the cell length and measured LifeAct-RFP intensity along a 10 pixel-thick line.

To quantify the enrichment of ECM in the mesoderm around PGCs at E7.5, the intensity of regions of the mesoderm containing PGCs were divided by the intensity of regions without PGCs using Fiji (Fig. 2b). To quantify the enrichment of ECM around individual PGCs (Fig. 2d, h, Ex-tended Data Fig. 2f-j, m-n), we traced 20 pixel-thick lines along the contours of PGCs and nearby somatic cells and measured the immunofluorescence intensity of laminin, collagen IV, fibronectin, or laminin reporters in Fiji. Measurements were normalized to the mean intensity around somatic cells in each sample to enable comparisons across samples.

To quantify changes in the basement membrane (Extended Data Fig. 3b-d), we performed two sets of measurements in Fiji and MatLab. First, intensity profiles along 20 pixel-thick lines were taken perpendicular to the basement membrane. These profiles were smoothed with a Savitzky-Golay filter and then the full width at half maximum was taken as an estimate of basement membrane thickness. Second, intensity profiles along 20-pixel-thick lines were taken by tracing along approximately 25 *µ*m of the basement membrane. These profiles were also smoothed with a Savitzky-Golay filter, the mean was subtracted, and then the variance was calculated as an estimate of basement membrane continuity. For both measurements, five profiles were obtained from different areas of the sample and averaged to obtain single values for each sample.

To quantify morphology and dynamics of PGCs entering and exiting the endoderm (Fig. 3), we analysed both fixed samples and live imaging data. For fixed samples, E7.5 embryos and E9.5 hindguts stained for laminin and the PGC marker Stella were examined to find PGCs spanning the basement membrane. As a proxy for constriction, we measured the width of the PGC at the level of the basement membrane in Fiji (Fig. 3b). For live imaging, we used tracking data obtained manually in MaMuT for PGCs and somatic cells from previously acquired light-sheet datasets[17]. We first defined the surface of the embryo by finding the convex hull of all somatic cell positions in a given time point, then determined the closest distance between each PGC and the outer surface of the convex hull using point2trimesh46. The results were plotted as a heatmap in RStudio, displaying only PGCs within 100 *µ*m of the surface. For protrusion measurements prior to PGC exit from the hindgut endoderm (Fig. 3h), we measured protrusion length and width (at the base of the protrusion) over time manually in Fiji. We then aligned these measurements for each PGC based on when the exit was completed, i.e. when the nucleus was no longer within the endoderm.

To quantify intercellular spaces (Fig. 4a, Extended Data Fig. 4b, f-g), we manually measured the widths of intercellular spaces based on 70 kDa dextran signal around PGCs and compared these across developmental stages and to manually measured PGC widths.

To quantify differences in DNA damage, we used three different commonly used markers, (*γ*H2AX, RPA32, and 53BP1), and quantified foci in PGCs and neighbouring somatic cells in the tissue of interest (Fig. 5b,d, Extended Data Fig. 5). For *γ*H2AX, we quantified the percentage of cells with >10 foci. For RPA32, we counted foci in each nucleus and reported the number of foci and the percentage of cells containing foci. For 53BP1, we counted foci in each nucleus and reported the number of foci and the percentage of cells containing ≥ 5 foci.

To quantify levels of lamins along the nuclear envelope (Fig. 5f, Extended Data Fig. 6c, d,f-g), we measured the immunofluorescence intensity of Lamin B1 and Lamin B2 in 10-pixel thick contours of nuclei generated based on DAPI. Intensity values were normalized to those of surrounding somatic cells within the tissue of interest to compare across samples. To quantify changes in the morphology of the nuclear envelope (Fig. 5f, Extended Data Fig. 6i), we counted indentations in the nuclear envelope based on immunostaining for the nuclear pore complex (MAB414 antibody).

#### AFM analysis

AFM force curves were analysed in MatLab. Force curves were smoothed using a Savitzky-Golay filter, then the first 250 nm of each curve were fit using the Hertz model,

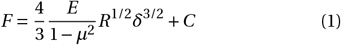

where F is the measured force, E is the elastic modulus, *µ* = 0.4 is the Poisson ratio[45], R is the radius of the probe (3.31 µm), *δ* is the tip-to-sample separation, and C is a constant.

## Supplementary Information

## Supplementary Data Figures

**Supplementary Figure 1:**
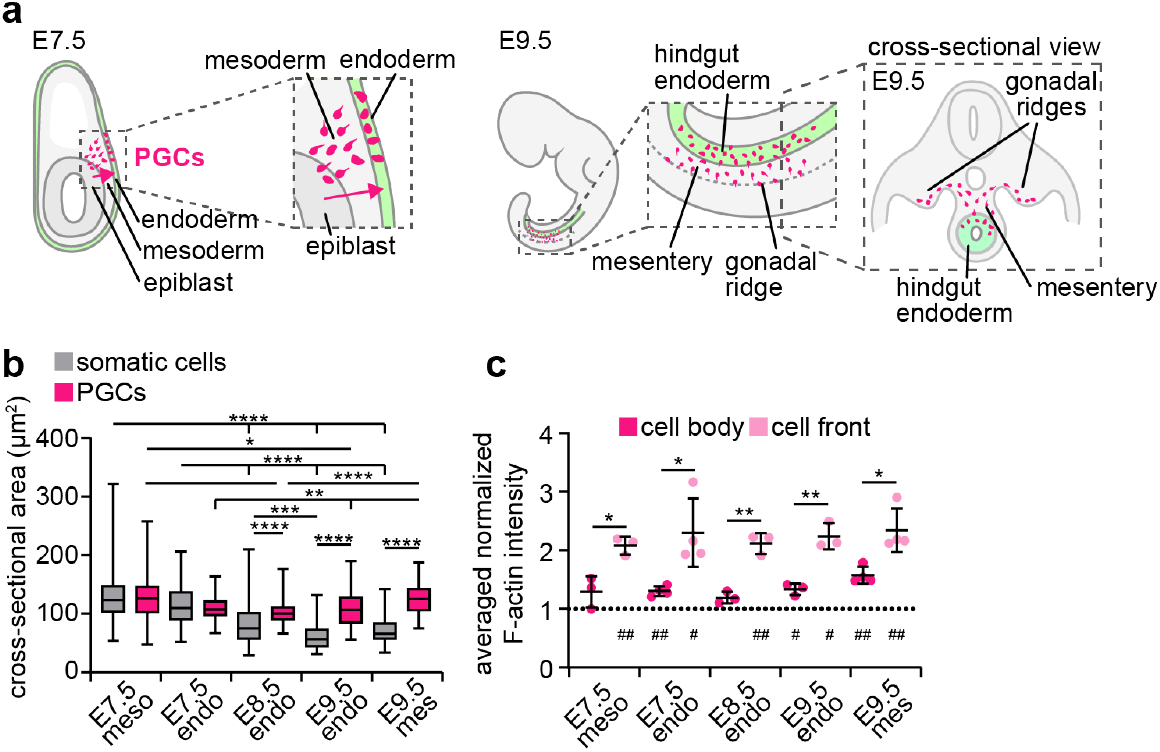
Cytoskeleton and cell adhesion in migrating PGCs. **a** Schematics showing zoomed-in views of regions that PGCs migrate through at E7.5 and E9.5. **b** Cross-sectional areas of PGCs and somatic cells from E7.5 to E9.5 in the mesoderm (meso), endoderm (endo), and mesentery (mes). **c** F-actin intensity at cell fronts and cell bodies normalized to neighbouring somatic cells in PGCs from E7.5 to E9.5 and in the mesoderm, endoderm, and mesentery. Averaged values for each embryo are shown. Asterisks are for pairwise comparisons using two-sided t-test, while symbols are for one-sample t-tests comparing data points to 1 (i.e., equal to somatic cell intensity) with indicating p <0.05, p<0.01.

**Supplementary Figure 2:**
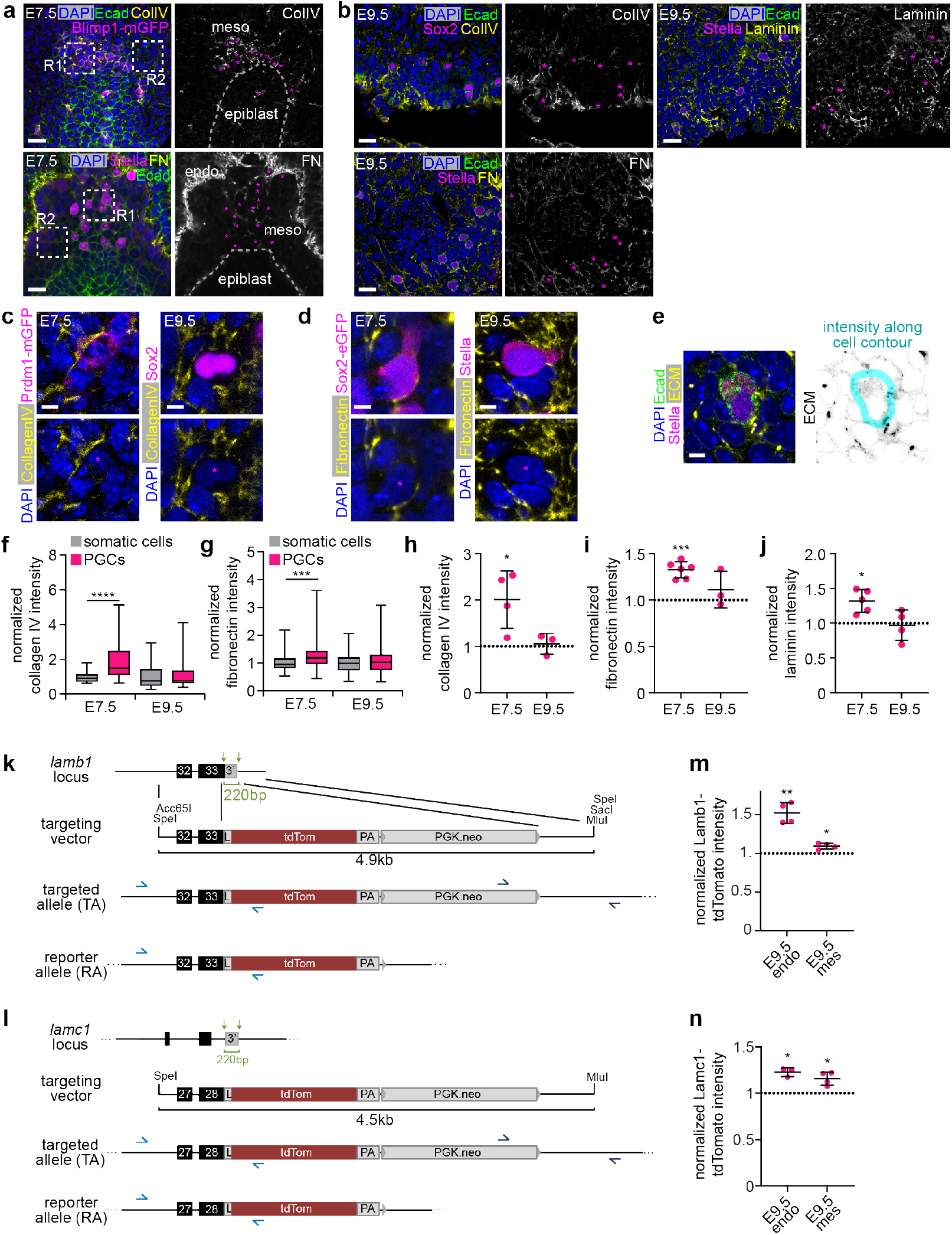
ECM around migrating PGCs. **a** Maximum intensity projections of the posterior regions of E7.5 embryos stained for the ECM components collagen IV or fibronectin, the PGC markers Stella or GFP in Sox2-eGFP embryos, Ecad, and DAPI. PGCs are indicated with magenta asterisks. **b** E9.5 hindguts stained for the ECM components laminin, collagen IV (ColIV) or fibronectin (FN), the PGC markers Stella or Sox2, Ecad, and DAPI. PGCs are indicated with magenta asterisks. Schematic indicates where images were taken. **c** PGCs in E7.5 embryos and E9.5 hindguts stained for collagen IV, GFP in Prdm1-mGFP embryos or Sox2 in WT, and DAPI. Magenta asterisks indicate PGCs. **d** PGCs in E7.5 embryos and E9.5 hindguts stained for fibronectin, GFP in Sox2-GFP embryos or Stella in WT, and DAPI. Magenta asterisks indicate PGCs. **e** Zoomed-in image of a PGC and the surrounding ECM. Contour indicates where ECM immunofluorescence intensity was measured. **f-g** Immunofluorescence intensity of collagen IV (f) and fibronectin (g) along PGC and somatic cell contours, normalized to the mean intensity around somatic cells in each embryo. **h-j** Collagen IV (h), fibronectin (i), and laminin (j) immunofluorescence intensity around PGCs normalized to somatic cells and averaged for each embryo at E7.5 and E9.5. **k-l** Targeting strategy using site directed homologous recombination for generation of Lamb1-tdTomato (k) and Lamc1-tdTomato (l) mice (see Methods; black: exons; 3’: endogenous 3’ UTR; PA: PolyA; L: linker; triangles: loxP sites; green arrows: TALEN binding sites; blue arrows: screening primers). **m-n** Lamb1-tdTomato (k) and Lamc1-tdTomato (l) live reporter intensity around PGCs normalized to somatic cells and averaged for each embryo. **** denotes p < 0.0001, *** p <0.001, ** p <0.01, * p < 0.05. Scale bars 25 *µ*m in (a-b), 5 *µ*m in (c-e).

**Supplementary Figure 3:**
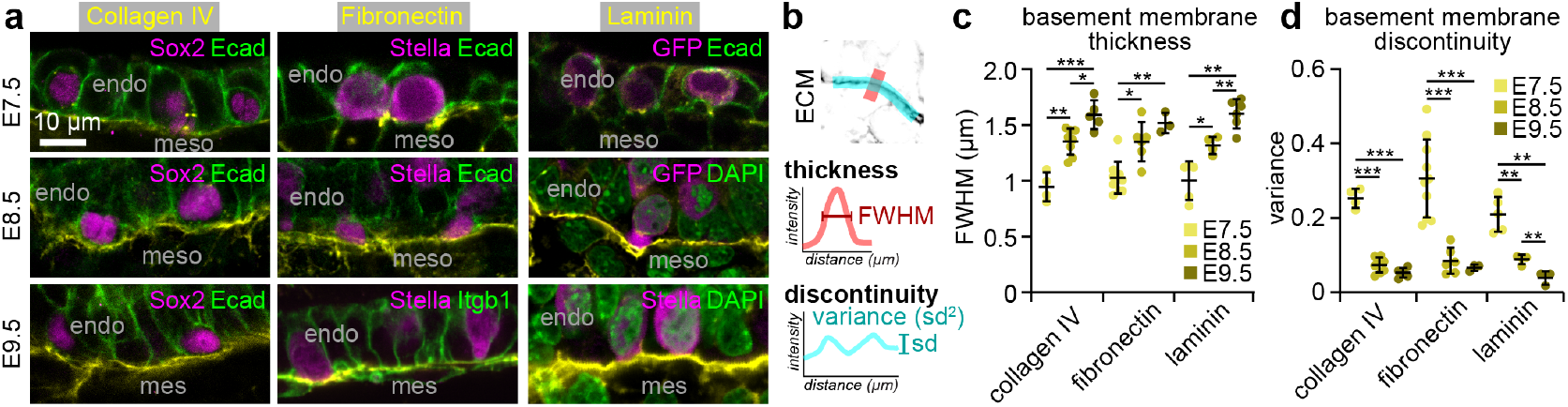
The endoderm basement membrane develops during PGC migration. **a** E7.5 and E8.5 embryos and E9.5 hindguts stained for the ECM components collagen IV, fibronectin, or laminin, the PGC markers Sox2, Stella, or GFP in Sox2-eGFP embryos, and either Ecad, integrin-1 (Itgb1), or DAPI to provide tissue context. Images are aligned such that the apical side of the endoderm is at the top and the basal side is at the bottom. **b**Schematics depicting measurements of basement membrane development. Full width at half maximum (FWHM) of perpendicular intensity profiles was used to estimate thickness. Variance of intensity profiles along the basement membrane was used to estimate discontinuity. **c** Thickness of the basement membrane at E7.5, E8.5, and E9.5. **d** Continuity of the basement membrane at E7.5, E8.5, and E9.5. ***denotes p <0.001, ** p <0.01, * p < 0.05. Scale bar 10 *µ*m.

**Supplementary Figure 4:**
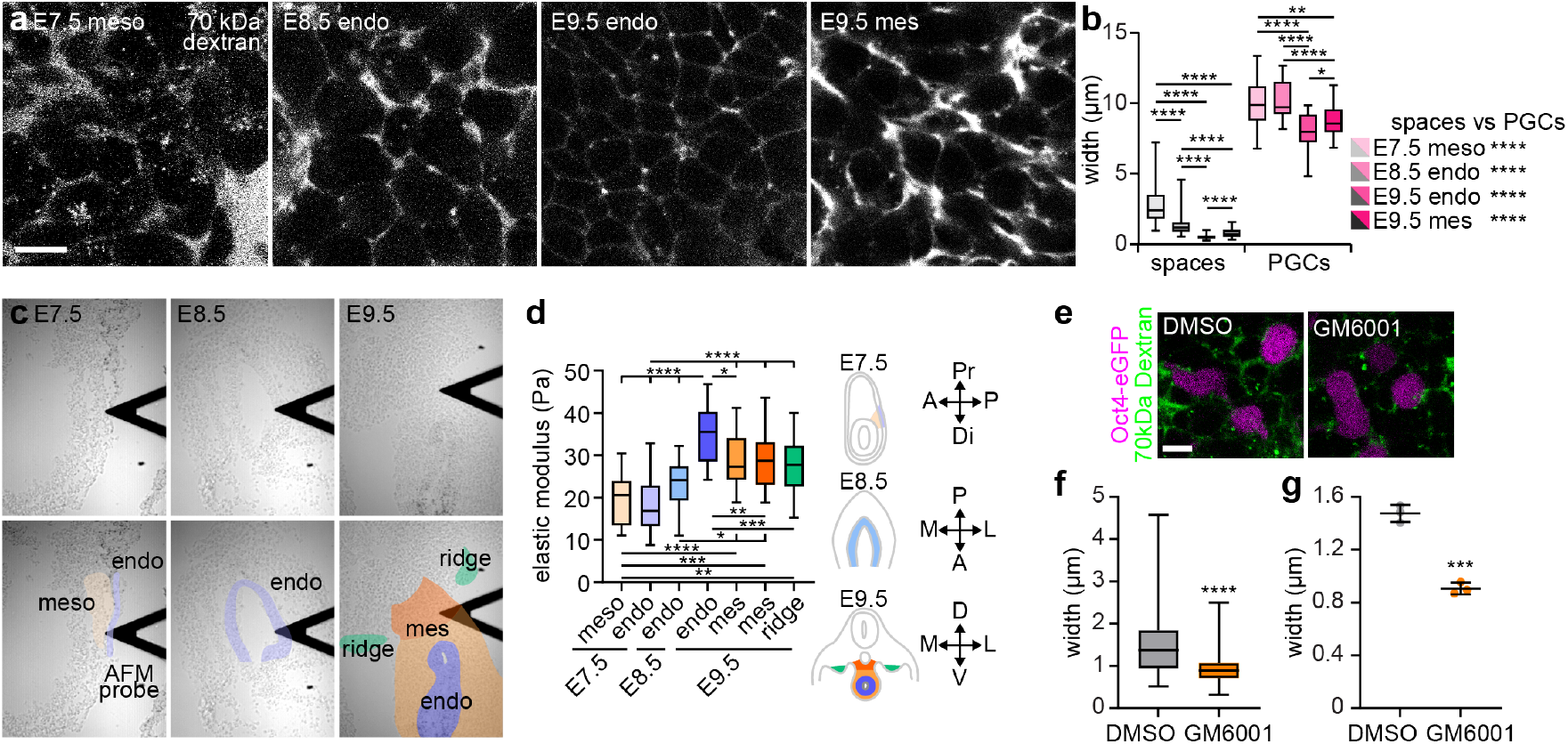
Changes in intercellular spaces and stiffness in tissues along PGC migratory path. **a** Snapshots of live E7.5 and E8.5 embryos and E9.5 hindgut incubated with 70kDa dextran. **b** Widths of all intercellular spaces measured (using Dextran signal) and all PGCs. **c** Brightfield images of tissue sections used for AFM, with the annotated version provided below the originals. **d** All measurements of elastic modulus of tissues around migrating PGCs from E7.5 to E9.5, as indicated by the schematics. **e** Hindguts expressing OCT4-eGFP treated with GM6001 or DMSO control and incubated with 70 kDa dextran to show intercellular spaces. **f** Widths of intercellular spaces in DMSO controls and GM6001-treated hindguts. g Mean widths of intercellular spaces in DMSO controls and GM6001-treated hindguts. **** denotes p < 0.0001, *** p <0.001, ** p <0.01, * p < 0.05. Scale bars 10 *µ*m.

**Supplementary Figure 5:**
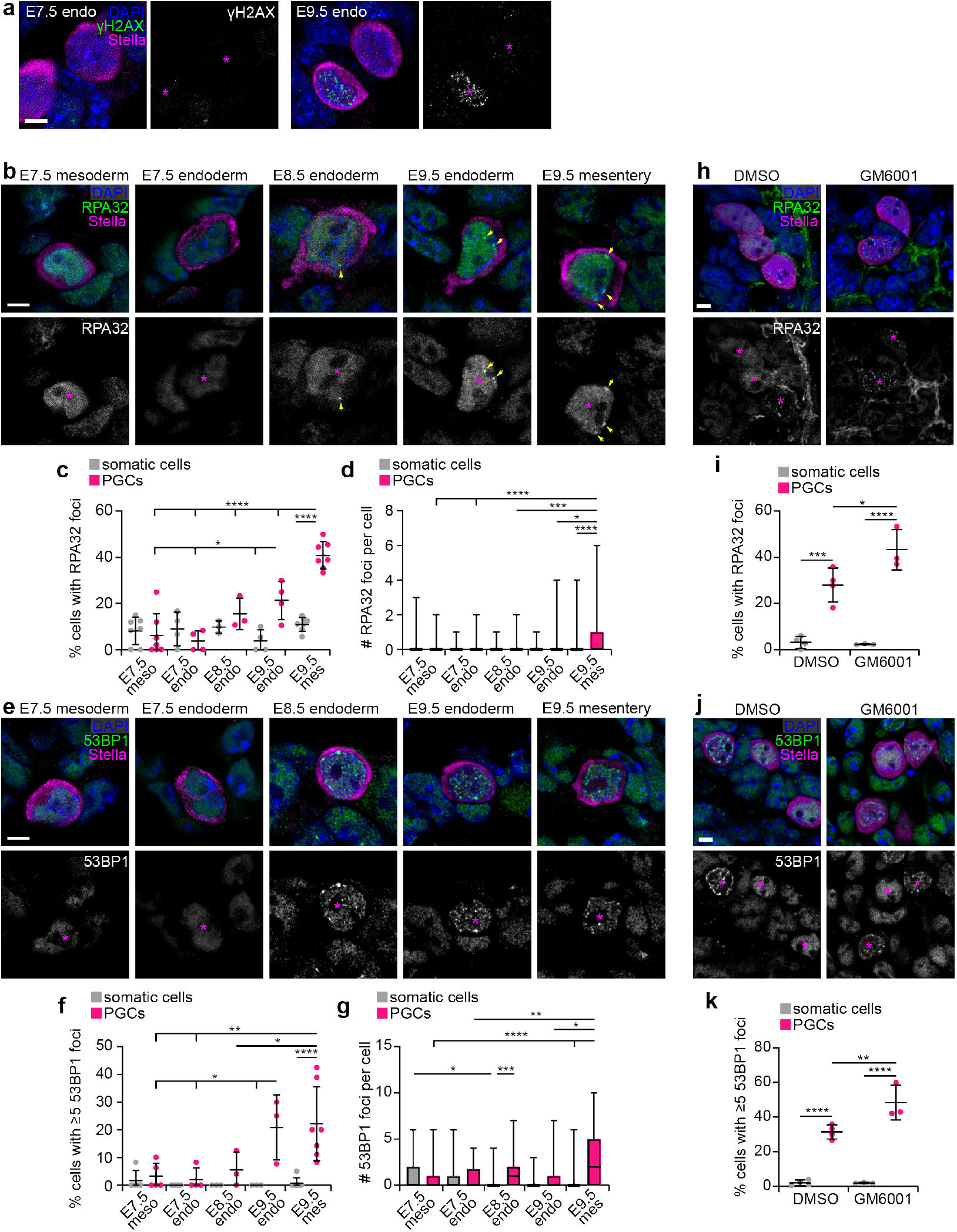
Migrating PGCs exhibit increasing evidence of DNA damage. **a** Sections of E7.5 and E9.5 embryos stained for *γ*H2AX, Stella and DAPI for tissues/stages not shown in Fig 5. Asterisks indicate PGCs. **b** Sections of E7.5, E8.5, and E9.5 embryos stained for the DNA damage marker RPA32 and the PGC marker Stella and counterstained with DAPI. PGCs are indicated with pink asterisks. Yellow arrows indicate RPA32 foci. **c** Percentage of cells with RPA32 foci in PGCs and in surrounding somatic cells in the relevant tissues at each developmental stage. **d** Number of RPA32 foci per cell in the mesoderm (meso), endoderm (endo), and mesentery (mes) at E7.5, E8.5, and E9.5. **e** Sections of E7.5, E8.5, and E9.5 embryos stained for the DNA damage marker 53BP1 and the PGC marker Stella and counterstained with DAPI. PGCs are indicated with pink asterisks. **f** Percentage of cells with ≥5 53BP1 foci in PGCs and in surrounding somatic cells in the relevant tissues at each developmental stage. **g** Number of 53BP1 foci per cell in the mesoderm, endoderm, and mesentery at E7.5, E8.5, and E9.5. **h** Sections of E9.5 hindguts cultured with DMSO or GM6001 and stained for RPA32, Stella and DAPI. Asterisks indicate PGCs. **i** Percentage of cells with RPA32 foci in PGCs and in surrounding somatic cells in E9.5 hindgut cultured with DMSO or GM6001. **j** Sections of E9.5 hindguts cultured with DMSO or GM6001 and stained for 53BP1, Stella and DAPI. Asterisks indicate PGCs. **k** Percentage of cells with ≥5 53BP1 foci in PGCs and in surrounding somatic cells in E9.5 hindgut cultured with DMSO or GM6001. **** denotes p < 0.0001, *** p <0.001, ** p <0.01, * p < 0.05. Scale bars 5 *µ*m.

**Supplementary Figure 6:**
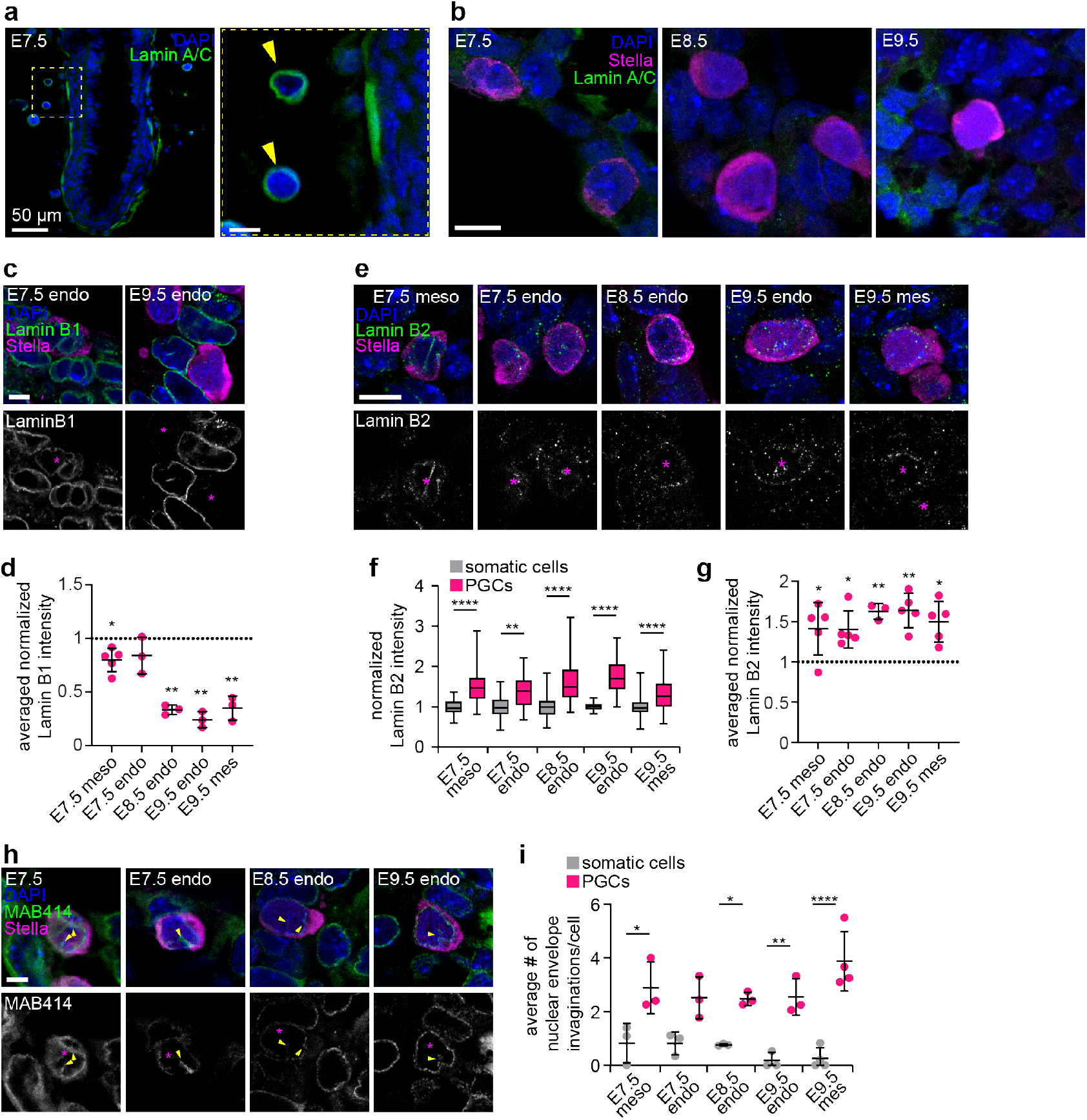
Nuclear lamina and nuclear morphology in migrating PGCs. **a** Section of an E7.5 embryo stained for Lamin A/C and DAPI. Lamin A/C is expressed in maternal decidual cells (yellow arrowheads), but not in embryonic cells. **b** Sections of E7.5, E8.5, and E9.5 embryos stained for Lamin A/C, Stella, and DAPI. **c** Sections of E7.5 and E9.5 embryos stained for Lamin B1, Stella and DAPI for tissues/stages not shown in Fig. 5. **d** Immunofluorescence intensity of Lamin B1 around the nucleus of PGCs in the relevant tissues at each developmental stage, normalized to the mean intensity of somatic cells in each sample and averaged for each embryo. **e** Sections of E7.5, E8.5, and E9.5 embryos stained for Lamin B2 and the PGC marker Stella and counterstained with DAPI. PGCs are indicated with pink asterisks. **f** Immunofluorescence intensity of Lamin B2 around the nucleus of PGCs and surrounding somatic cells in the relevant tissues at each developmental stage, normalized to the mean intensity of somatic cells in each sample. **g** Immunofluorescence intensity of Lamin B2 around the nucleus of PGCs in the relevant tissues at each developmental stage, normalized to the mean intensity of somatic cells in each sample. **h** Sections of E7.5, E8.5, and E9.5 embryos stained for MAB414, Stella and DAPI for tissues/stages not shown in Fig. 5. Yellow arrows indicate invaginations of the nuclear envelope. **i** Average number of nuclear envelope invaginations per cell. **** denotes p < 0.0001, ** p <0.01, and * p <0.05. Scale bars 10 µm unless otherwise indicated.

## Movies

**Movie 1** Protrusive migration of a PGC in the mesentery of an E9.5 hindgut explant expressing Sox2-eGFP and mKate2-nls.

**Movie 2** F-actin-rich protrusion formation by a PGC in the hindgut endoderm of an E9.5 hindgut explant expressing Sox2-eGFP and LifeAct-RFP.

**Movie 3** PGC migration coincides with the appearance of laminin in an E7.5 embryo expressing Sox2-eGFP and Lamb1-tdTomato.

**Movie 4** Dynamic and long-lived protrusion into the mesentery made by a PGC in the hindgut endoderm of an E9.5 hindgut explant expressing Oct4-eGFP and Lamc1-tdTomato.

**Movie 5** Protrusion extension, widening, and PGC exit from the hindgut endoderm in an E9.5 hindgut explant expressing Sox2-eGFP and mKate2-nls.

**Movie 6** PGC exiting the hindgut endoderm and immediately fragmenting in an E9.5 hindgut explant expressing Sox2-eGFP and mKate2-nls.

**Movie 7** DMSO control and GM6001-treated hindgut explants expressing Sox2-eGFP showing normal migration in controls and fast migration followed by cell rupture when MMP activity is inhibited and confinement is increased.

## Supplementary Data Table 1

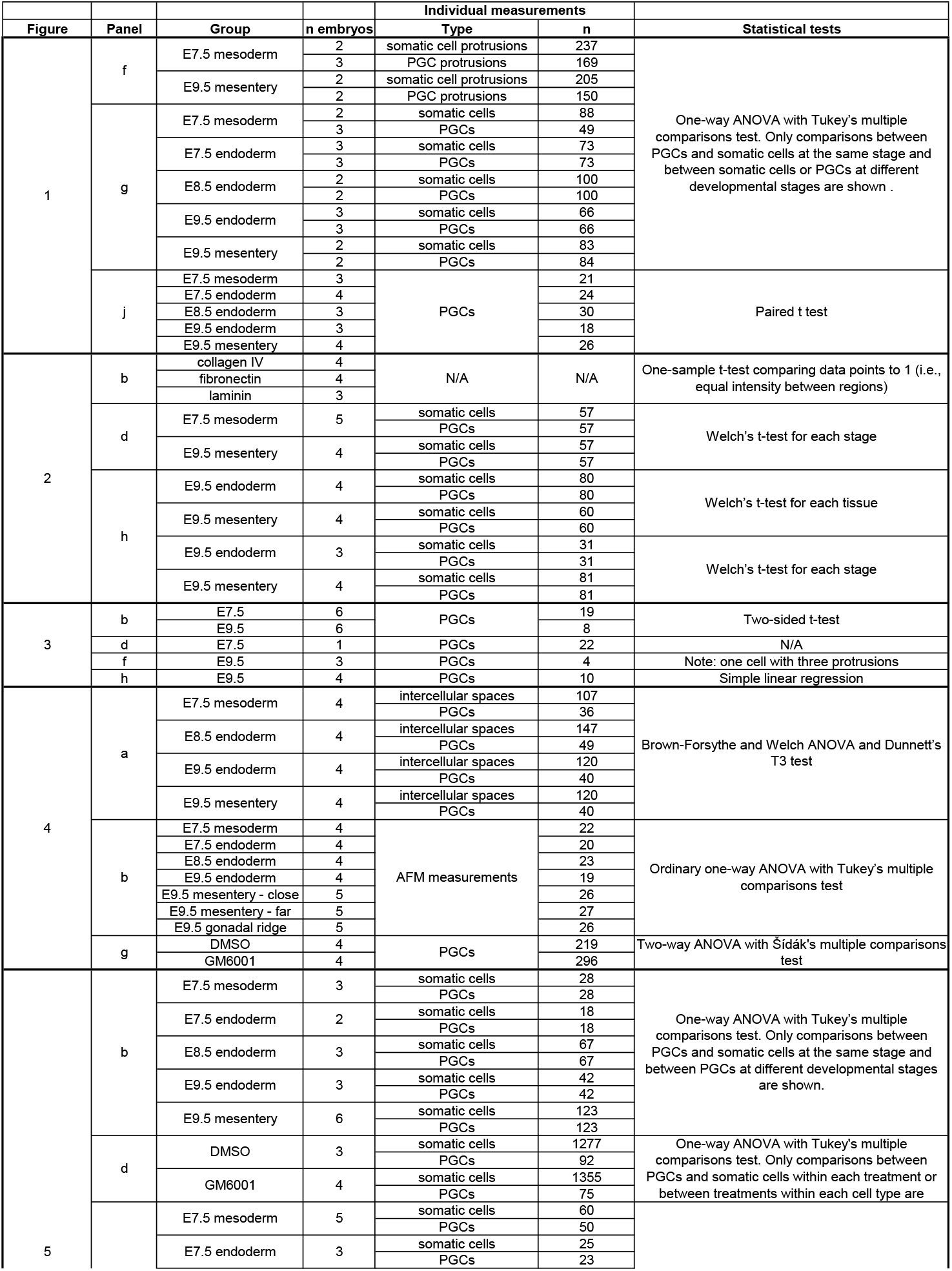
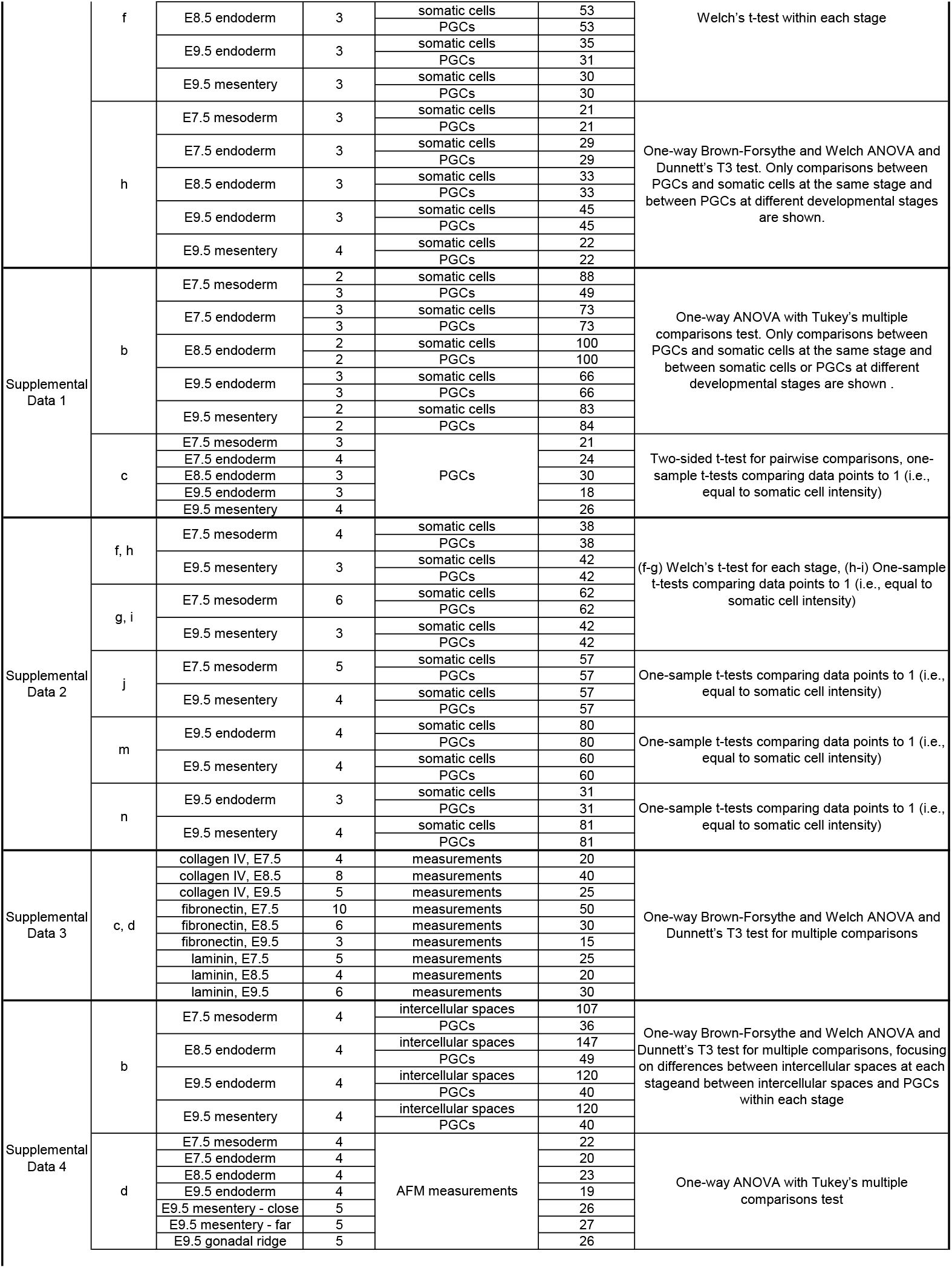
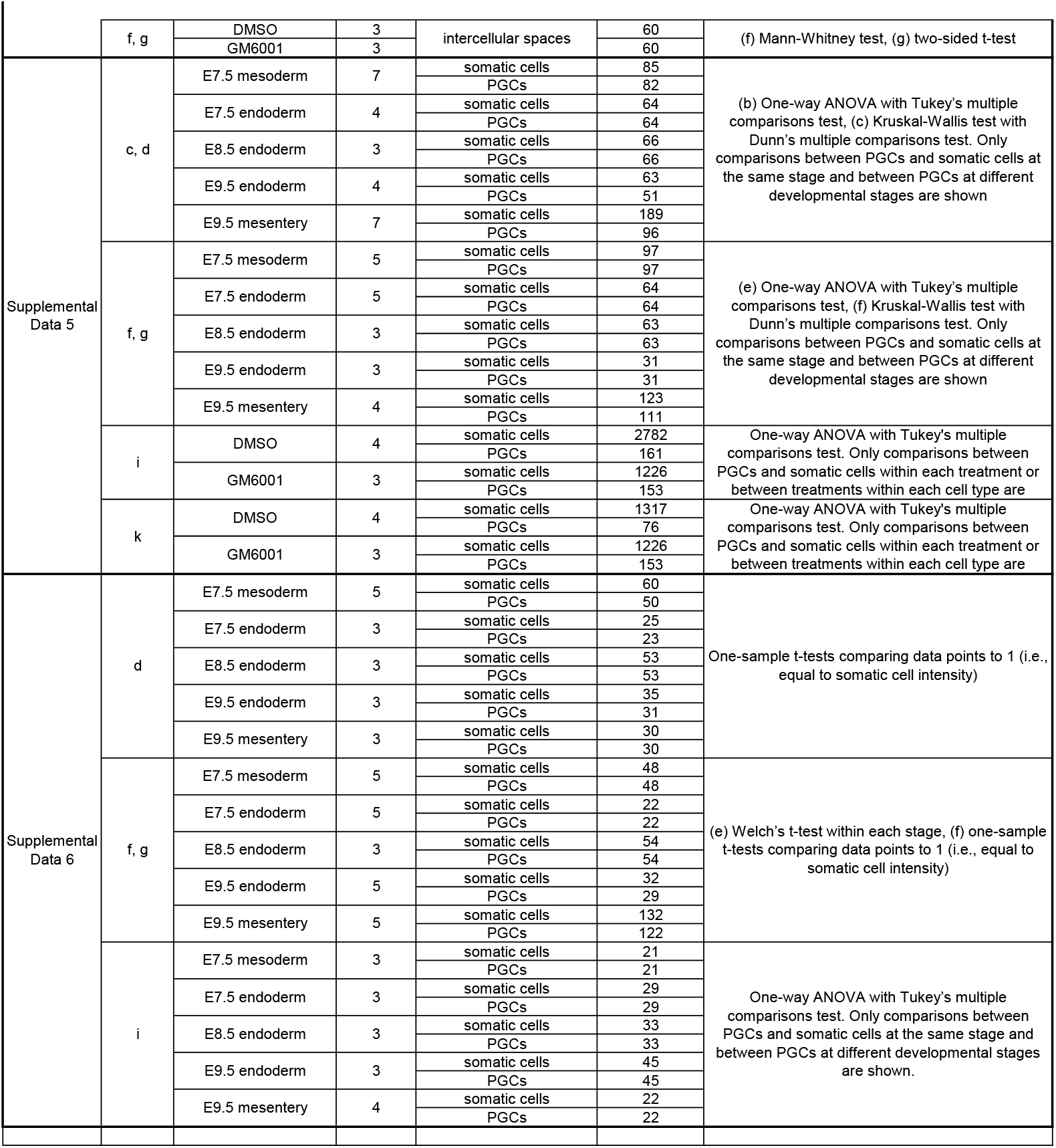

